# Improved *in situ* Characterization of Proteome-wide Protein Complex Dynamics with Thermal Proximity Co-Aggregation

**DOI:** 10.1101/2023.02.13.528386

**Authors:** Siyuan Sun, Zhenxiang Zheng, Jun Wang, Fengming Li, An He, Ruijun Tian, Chris Soon Heng Tan

## Abstract

Vast majority of cellular activities are carried out by protein complexes that assembled dynamically in response to cellular needs and environmental cues. Large scale efforts had uncovered a large repertoire of functionally uncharacterized protein complexes which necessitate new strategies to delineate their roles in various cellular activities and diseases. Thermal proximity co-aggregation profiling could be readily deployed to simultaneously characterize the dynamics for hundreds to thousands of protein complexes *in situ* across different cellular conditions. Toward this goal, we had optimized the original method both experimentally and computationally. In this new iteration termed Slim-TPCA, fewer temperatures are used which increase throughputs by over 3X, while coupled with new scoring metrics and statistical evaluation resulted in minimal compromise in coverage and the detection of more relevant protein complexes. Overall, less samples are needed, false positives from batch effects are minimized and statistical evaluation time is reduced by two orders of magnitude. We applied Slim-TPCA to profile state of protein complexes in K562 cells under different duration of glucose deprivation. More protein complexes are found dissociated based on TPCA signature in accordance with expected downregulation of most cellular activities. These complexes include 55S ribosome and various respiratory complexes in mitochondria revealing the utility of TPCA to study protein complexes in organelles. On other hand, protein complexes involved in protein transport and degradation are found increasingly associated revealing their involvement in metabolic reprogramming during glucose deprivation. In summary. Slim-TPCA is an efficient strategy for proteome-wide characterization of protein complexes. The various algorithmic improvement of Slim-TPCA is available as Python package at https://pypi.org/project/Slim-TPCA/

## Introduction

Proteins are molecular workforces performing virtually all biological processes in cells but they seldom act alone. Instead, many are part of higher-order molecular machines commonly termed protein complexes that are assembled when needed from protein-protein interactions (PPIs)^2^. The interactions among proteins are often dynamic and regulated by various mechanisms including temporal posttranslational modifications and spatial redistribution of proteins that changes protein functionalities^3^. The study of PPIs can uncover the underlying machineries of various cellular processes and reveal how their dysregulation could lead to diseases, thus providing insight for development of new therapeutic treatments including the identification of new drug targets^4^. As such, identifying and characterizing the dynamics of PPIs and protein complexes are among the endeavors of biological sciences with increasing attention and research efforts in recent years^5-7^.

Experimental methods for studying PPIs can be broadly classified into traditional hypothesis-driven methods (e.g. immunoprecipitation, yeast two-hybrid) and high-throughput methods. Many of the latter are coupled with protein mass spectrometry (MS) for protein identification which include co-fractionation/elution and crosslinking of protein complexes^8,9^. In addition, a variety of computational models have also been proposed for predicting PPIs^10^. While experimental methods have their own limitations, they are often hailed as the gold standard used in the validation of computational approaches. Traditional experimental methods such as Y2H^11^, LUMIER^12^ and AP-MS^5^ can be scaled up to obtain large sets of PPI data but often suffer from higher false-negative rate that necessitated a combination of complementary experimental methods to furnish a more comprehensive interactome. However, these methods require either protein engineering (e.g., epitope tagging) or the use of antibodies specific to target proteins, and are not applicable *in situ*.^13^. Importantly, current large-scale interactome profiling are carried out with specific cell lines under basal conditions, thus the conservation and dynamics of identified PPIs and protein complexes across cell lines, cellular states and physiological conditions are unclear.

In 2018, we conceived the Thermal Proximity Co-Aggregation (TPCA) method, pioneering the use of thermal proteome profiling to study the proteome-wide dynamics of PPIs^14^ and protein complexes *in situ*. The underlying experimental method of TPCA profiling is similar to the TPP/MS-CETSA method where proteins are subjected to a gradient of denaturing temperature. The solubility of protein decreases with increasing temperature which is quantified with mass spectrometry^15,16^, resulting in a “melting curve” that are often influenced by protein-ligand interactions. TPP/MS-CETSA permits the measurement of protein-ligand binding in intact cells and is widely used in drug target deconvolution^17^ and off-target studies^18^. Our previous work revealed that signature of PPIs is embedded in thermal proteomics data^14^ as similar thermal melting curves of co-aggregating or co-precipitating proteins in the same protein complex. Arguably, TPCA profiling is one of the few methods that provide a proteome-wide approach to detect PPIs occurring in intact cellular environment^9,19^.

The TPCA method for analyzing protein-protein interactions has been independently demonstrated and applied since its conception. For example, the principle of TPCA was applied to study the modulation of protein complexes across different phases of cell cycle^20,21^. The complexes that are regulated in different phase of cell cycle were shown to have high correlation with the corresponding biological activities. Hashimoto et al. further applied TPCA to profile the dynamics of host protein complexes in different stages of viral infection and demonstrated that the TPCA can used to study PPIs between proteins from different species^1^. Moreover, TPCA had been demonstrated in thermal proteomic data obtained from tissue and blood samples^22^ where subunits of protein complexes are grouped hierarchically based on their median melting temperature (*T*_*m*_) to assess their assembly state across different tissues. In 2021, Kalxdorf et al. used the TPCA method to analyze complex formation during T-cell activation using changes in the distance between attachment proteins and the core of complexes^23^. To facilitate the use of TPCA for profiling PPIs, the computational workflow for TPCA analysis has been integrated into rTPCA^24^ and ProSAP^25^ software analysis packages.

Current TPCA workflow provides statistical evidence for the modulation of protein complexes using existing databases of protein complexes as reference^26,27^. In recent years, the repertoire of protein complex have expanded greatly with large scale interactome profiling projects^5-7^ and the development of system-level complex profiling techniques^19^ including co-fractionation-MS^28-30^ and protein network-based clustering algorithms^31,32^. However, the conservation and functions of these putative complexes across different cell types and cellular states are unclear.

Conceptually, without the need for antibodies and epitope-tagging of proteins, the TPCA method can be readily deployed to rapidly characterize the dynamics of multiple uncharacterized protein complexes simultaneously under different cellular conditions and perturbations. This could aid in the function annotation of these complexes. Nonetheless, there are several aspects where the TPCA method can be further optimized. In the original TPCA method, 10 samples are heated to different temperatures in gradient followed by abundance quantification of soluble protein using protein mass spectrometry. The difference in thermal solubility between proteins is measured by Euclidean distance, and the significance is estimated by a bootstrap sampling algorithm^14^. Thus, large number of sample and long computational time are needed in conventional TPCA method. Furthermore, one set of isobaric tandem mass tags (TMT) reagent with 10 channels is used to label only one set of TPCA experiment for a condition, thus the control and experimental samples had to be analyzed by mass spectrometry at different time with batch effect that contribute to deviation in measurements^33^.

Here, we optimize the TPCA method and propose a new streamlined version called Slim-TPCA. Slim-TPCA was optimized in both algorithmic and experimental aspects. First, in Slim-TPCA protocol, fewer temperature points are used which provide more flexibility in experimental design and help reduce batch effect (Fig.1a). Next, through algorithm optimization, Slim-TPCA can better identify PPIs and dynamic changes in protein complexes with improved statistical power. We applied the optimized Slim-TPCA method to analyze the dynamic changes of protein complexes at five different time points of glucose deprivation in the same MS analysis. We successfully identified many protein complexes known to associate with glucose starvation and revealed new connections with many other complexes. Slim TPCA is available as an independent python package to facilitate analysis of protein complexes.

**Fig. 1.**
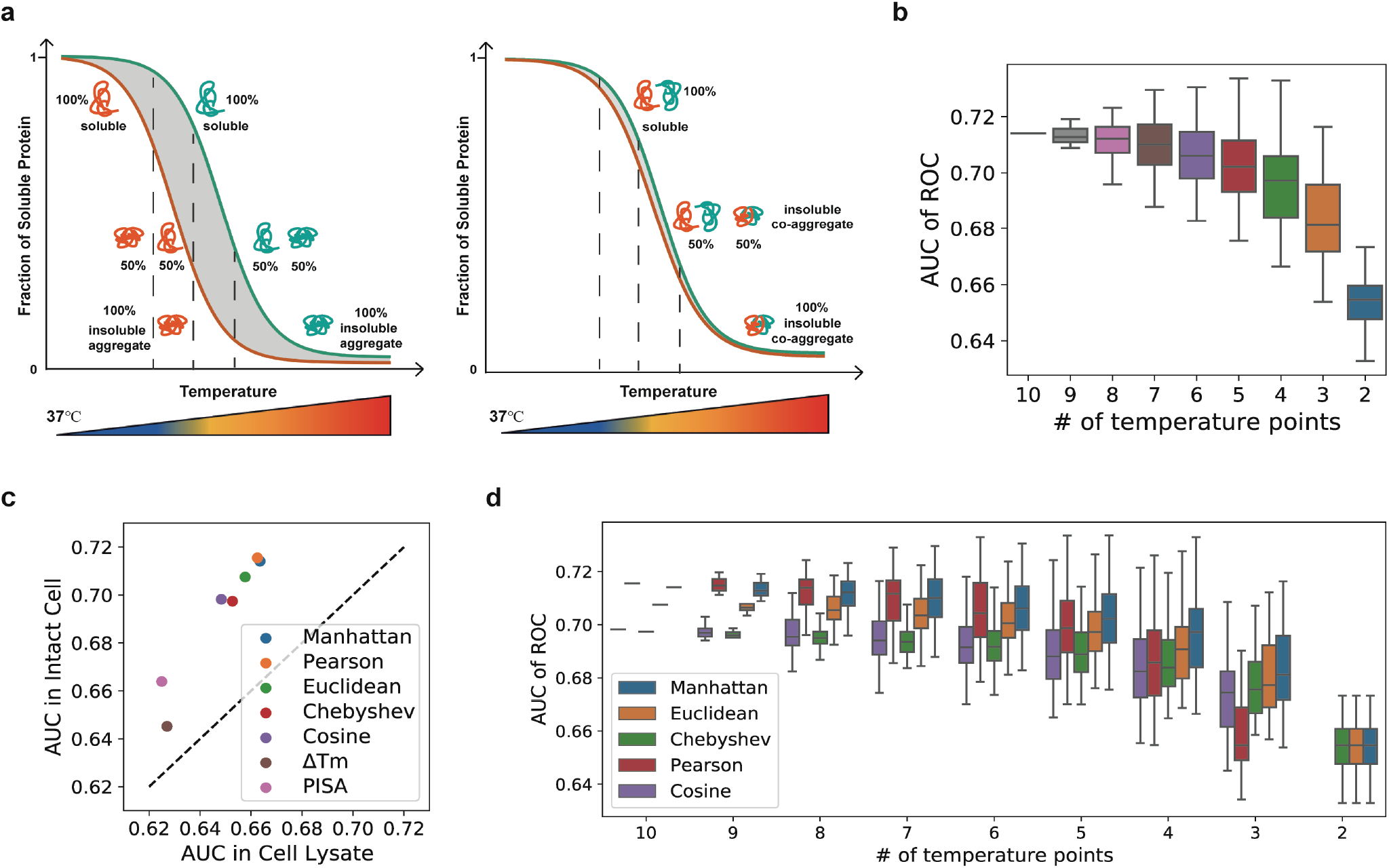
Distance measurements evaluated in Slim-TPCA. **(a)** Principle of Slim-TPCA for monitoring protein-protein interactions based on similarity between protein melting curves. In Slim-TPCA, lesser evenly spaced temperature points is used instead of the 10 gradient temperature points in the conventional method. **(b)** Predictive power of TPCA signature with different number of temperature points for differentiating between interacting and non-interacting protein pairs. All combinations of temperature points are tested and predictive power generally decreases with less temperature points. **(c)** Predictive power of TPCA signature quantified by different measures across 10 temperature points for differentiating between interacting and non-interacting protein pairs. **(d)** The ability of different measures in predicting PPI when used with fewer temperature points. All combinations of temperature were tested accordingly. The Pearson correlation coefficient is not applicable to Slim-TPCA with fewer temperature points, while the Manhattan distance retains good detection ability when using 3 temperature points.

## Results

### Fewer temperature points in Slim-TPCA

The fundamental basis of TPCA profiling for characterizing protein-protein interactions and their dynamics is the similarity in their melting curves quantified as solubility similarity of interacting proteins across multiple denaturing temperatures. In the first version of TPCA method, samples are subjected to 10 gradient temperatures, followed by isobaric TMT labelling and MS analysis to infer the so-called “melting” curve of proteins. Such a workflow requires a large number of samples, entailing multiple set of TMT reagents to label and analyze samples from different conditions (e.g., drug group and control group with replicates), which can lead to batch variation in MS measurement. As such, we seek to modify the original TPCA approach so that fewer temperatures can be used to analyze PPIs with minimal loss in information. We first assess the feasibility of doing so using thermal solubility data of K562 generated from cell lysate and intact cell that were published in the proof-of-concept paper^14^. The data is used to evaluate the TPCA signature between proteins for differentiating known PPIs from random protein pairs, and the predictive power is quantified with Area Under the Curve (AUC) of the Receiver Operating Characteristic (ROC) curve. In a nutshell, high AUC value indicates higher predictive (differentiating) power.

First, we analyzed the predictive power of different number of temperature points by testing all combination of temperatures with Euclidean distance measure used in the original work (Fig.1c). As expected, the AUC of ROC decreases with fewer temperature points but the decrease is surprisingly gradual. For example, the median AUC of two, three and four temperature points (including mandatory 37°C) is 0.65, 0.68 and 0.69 compared to AUC of 0.71 with 10 temperature points. Thus, we conclude that TPCA analysis can be performed using fewer temperature points with slight loss in predictive power.

Next, we analyzed which distance/similarity measures are most suitable for use with fewer temperature points. We analyzed 6 additional distance/similarity measures namely: Manhattan distance, Chebyshev distance, Cosine distance, Pearson’s correlation coefficient, PISA (area under melting curves), and Δ*T*_*m*_(melting temperatures). We observed the AUC values using 10 temperature point data with Manhattan distance and Pearson’s correlation coefficient are higher than that using Euclidean distance for both cell lysate and intact cell data (Fig.1c and Supplementary Fig.1). However, the performance of Pearson’s correlation coefficient deteriorates rapidly with fewer temperature points (Fig.1d) compared to using Manhattan distance and Euclidean distance. Overall, Manhattan distance gives the best performance across different number of temperature points and is used in our subsequent **-** analysis.

### Temperature points selection in Slim-TPCA

Next, we identified the combination of 2 and 3 temperature points (excluding 37°C) that produce the highest correlation with 10 temperature points for random protein pairs using Manhattan distance. Here, we assume that better correlation indicates higher coherency in the TPCA signature embedded in the data. Our analysis revealed that 37°C, 49°C and 58°C are the most suitable for 3 temperature points combination (Fig.2a and Supplementary Fig.2), while 37°C, 46°C, 55°C and 61°C are the most suitable for 4 temperature points combination (Fig.2b). We then used the identified combination of temperature points to analyze another set of TPCA data obtained from K562 cells which were treated with methotrexate (MTX). Previously, we performed bootstrapping to estimate the statistical significance of TPCA signature (non-random similarity among melting curves of proteins) observed in CORUM complexes. Here, we performed similar analysis using Manhattan distance comparing the TPCA signatures derived with 3 and 4 temperature points versus 10 temperature points. Encouragingly, the statistical significance of TPCA signatures of the complexes in the CORUM database calculated using our selected 3 and 4 temperature points are highly similar to those calculated at 10 temperature points with Pearson’s *r* greater than 0.93 and 0.96 for 3 and 4 temperature points respectively (Fig.2c). Next, we assessed the ability of the identified combination of temperature points to identify protein complexes dynamically modulated by MTX treatment. Similar to previous work, the statistical significance of difference in TPCA signature (termed dynamic *p*-value) across two conditions was estimated with bootstrapping analysis. We also observed high correlation in statistical significance estimated with 10 temperature points to that estimated with 3 and 4 temperature points with Pearson’s *r* greater than 0.92 and 0.94 respectively (Fig.2d and Supplementary Fig.3). Thus, our identified combinations of 3 or 4 temperature points are in good agreement with those calculated at 10 temperature points.

**Fig. 2.**
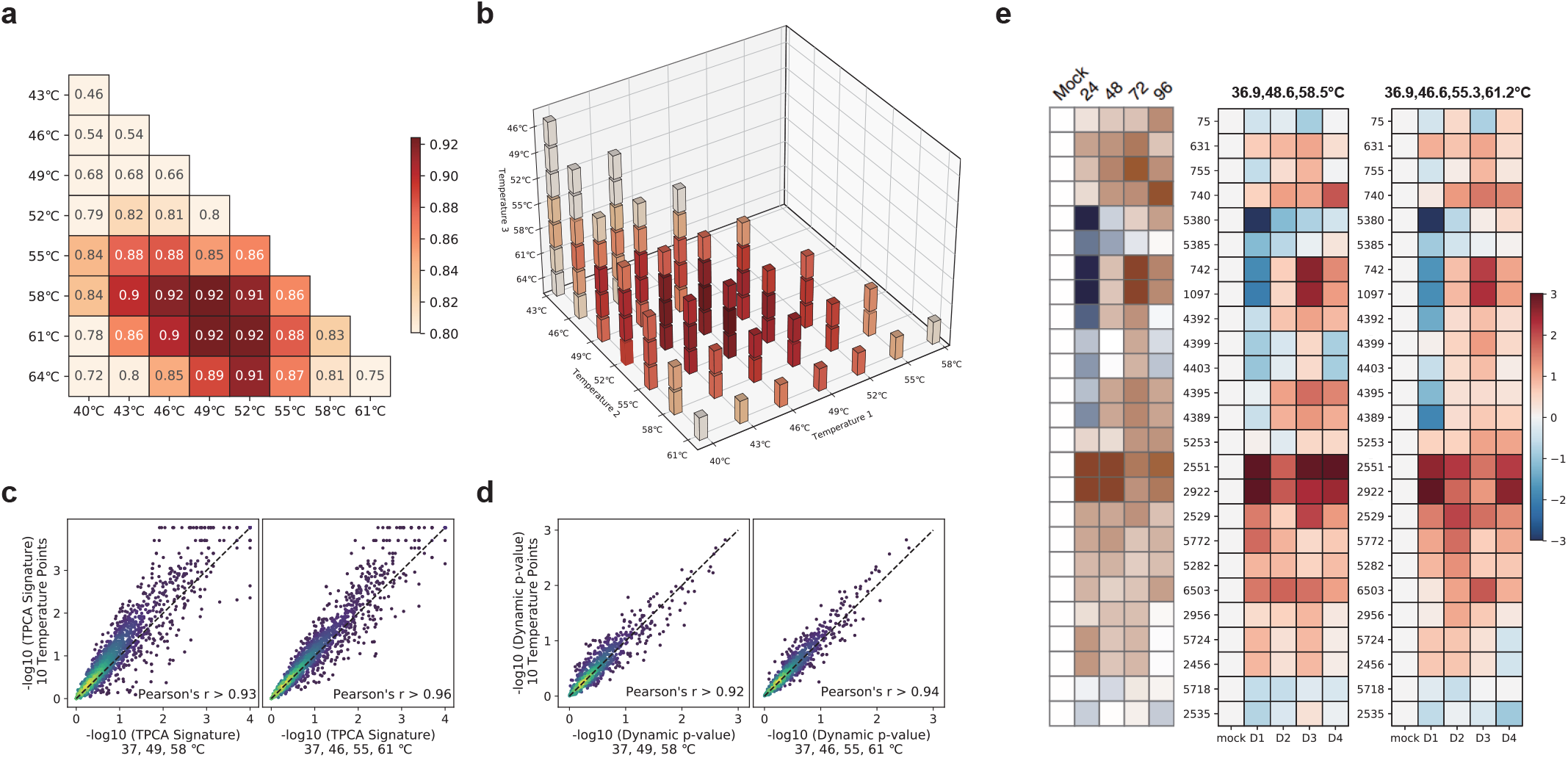
Selection of optimal combinations of temperature points for TPCA profiling. **(a)** Correlation of Manhattan distances for 100,000 random protein pairs calculated using 3 temperature points and 10 temperature points. The combinations of 3 temperature points ∈ {37°C, T_a_°C, T_b_°C}, where T_a_, T_b_∈ {40°C, 43°C, 46°C, 49°C, 52°C, 55°C, 58°C, 61°C, 64°C}. **(b)** Correlation of Manhattan distances for 100,000 random protein pairs calculated using 4 temperature points with those calculated using 10 temperature points. The combinations of 4 temperature points ∈ {37°C, T_a_°C, T_b_°C, T_c_°C}, where T_a_, T_b_, T_c_∈ {40°C, 43°C, 46°C, 49°C, 52°C, 55°C, 58°C, 61°C, 64°C}. **(c)** Statistical significance of TPCA signature for CORUM complexes (p-value) quantified with 3 and 4 temperature points as compared to 10 temperature points. **(d)** Statistical significance of dynamic (modulated) TPCA signatures for CORUM complexes (dynamic p-value) quantified with 3 and 4 temperature points as compared to 10 temperature points. **(e)** Variation in TPCA signatures for identified modulated protein complexes during virus infection. The statistical significance obtained with 10 temperature points (left^1^) can basically be recapitulated using 3 temperature points (middle, 36.9°C, 48.6°C, and 58.5°C) and 4 temperature points (right, 36.9°C, 46.6°C, 55.3°C, and 61.2°C). The figure for 10 temperature points is as obtained from the original publication^1^ where the color scheme for the 3 and 4 temperature points are matched as close as possible.

To further investigate the coherence of the identified combinations of temperature points across different experimental data, we used the data published in Hashimoto et al.^1^ to validate the selection of temperature points. In the work, the authors applied TPCA profiling to study the dynamics of protein complexes and identified a subset that are modulated during human cytomegalovirus (HCMV) infection.

Using Manhattan distance and similar sets of temperature points, the dynamics of identified modulated complexes across 4 days of infection obtained by 10 temperature points (Fig.2e, left panel) are generally consistent with the results calculated using 3 temperature points of 36.9°C, 48.6°C, 58.5°C (Fig.2e, middle panel) and 4 temperature points of 36.9°C, 46.6°C, 55.3°C, and 61.2°C (Fig.2e, right panel) This demonstrates that the identified combinations of temperature points are generally applicable across samples from different experiments.

Importantly, the identified optimal combination of temperature points is in line with our expectation that picking the intermediate, widely separated temperature points will maximally encapsulated TPCA signal embedded across temperature range tested. Using Slim-TPCA with 3 or 4 temperature points, we can now analyze samples under different conditions or multiple sets of experimental conditions, such as different treatment duration using a set of TMT reagents. As such, Slim-TPCA has the advantage of being able to reduce sample consumption, reduce batch effects, and provide more flexibility in experimental design.

### Relative distance algorithm optimization to detect dynamic complexes

The TPCA method has the capacity to identify protein complexes modulated across conditions based on changes in TPCA signature. Specifically, to detect dynamically modulated protein complexes, the average difference in solubilities between subunits in a protein complex are compared across conditions and benchmark against 10,000 randomly generated protein sets of the same size. Protein complexes that experience more extreme variations than artificially generated protein sets are considered as dynamically modulated protein complexes, which is assessed with the dynamic TPCA p-values. In the original TPCA method, the variation is obtained by direct subtraction of their average distances under different conditions, which is we termed as the absolute distance algorithm. This absolute distance algorithm has been demonstrated to detect known activated complexes, such as the CAF-1 complex formed at replication fork under MTX treatment which trapped K562 cells in S-phase of cell cycle. Nevertheless, the absolute distance algorithm can miss modulated complexes that already has strong TPCA signature in a reference condition but are further strengthen in another condition. For example, we will expect a population of cells to be in different phase of cell cycle with associated protein complexes exhibiting certain degree of TPCA signature. While the stoichiometry of such complexes could further peak, changes in the absolute distance are limited given the relatively strong TPCA signature of these complexes at basal conditions. In addition, proteins could also have similar thermal solubility independent of each other that are enhanced by interaction.

To detect protein complexes in above-mentioned situations, we introduce a new scoring algorithm based on relative distance to identify such complexes (see Methods). Specifically, changes in TPCA signature are quantified as a ratio of basal TPCA signature. We evaluate this relative distance algorithm using the sets of data from previous work where K562 cells enriched in S-phase of cell cycle using MTX are compared to non-synchronized cells. First, using the same method of sampling to generate random protein sets, we analyzed the correlation between the dynamic TPCA p-values obtained by the relative distance algorithm and the absolute distance algorithm. The dynamic TPCA signatures (p-values) of CORUM complexes calculated using the relative distance algorithm maintain good consistency with those calculated using the absolute distance algorithm for both 10-temperature TPCA and 4-temperature Slim-TPCA method (Fig.3a-b). Importantly, insignificant (high) p-values are highly consistency across absolute and relative distance algorithm but the latter results in enhanced p-value for a subset of complexes that already have low p-value from absolute distance scoring. In other words, the relative distance algorithm neither lead to depreciation of p-value estimated nor enhancement for those with very insignificant p-value but identified a subset of complexes with enhanced p-value indicating increased sensitivity.

**Fig. 3.**
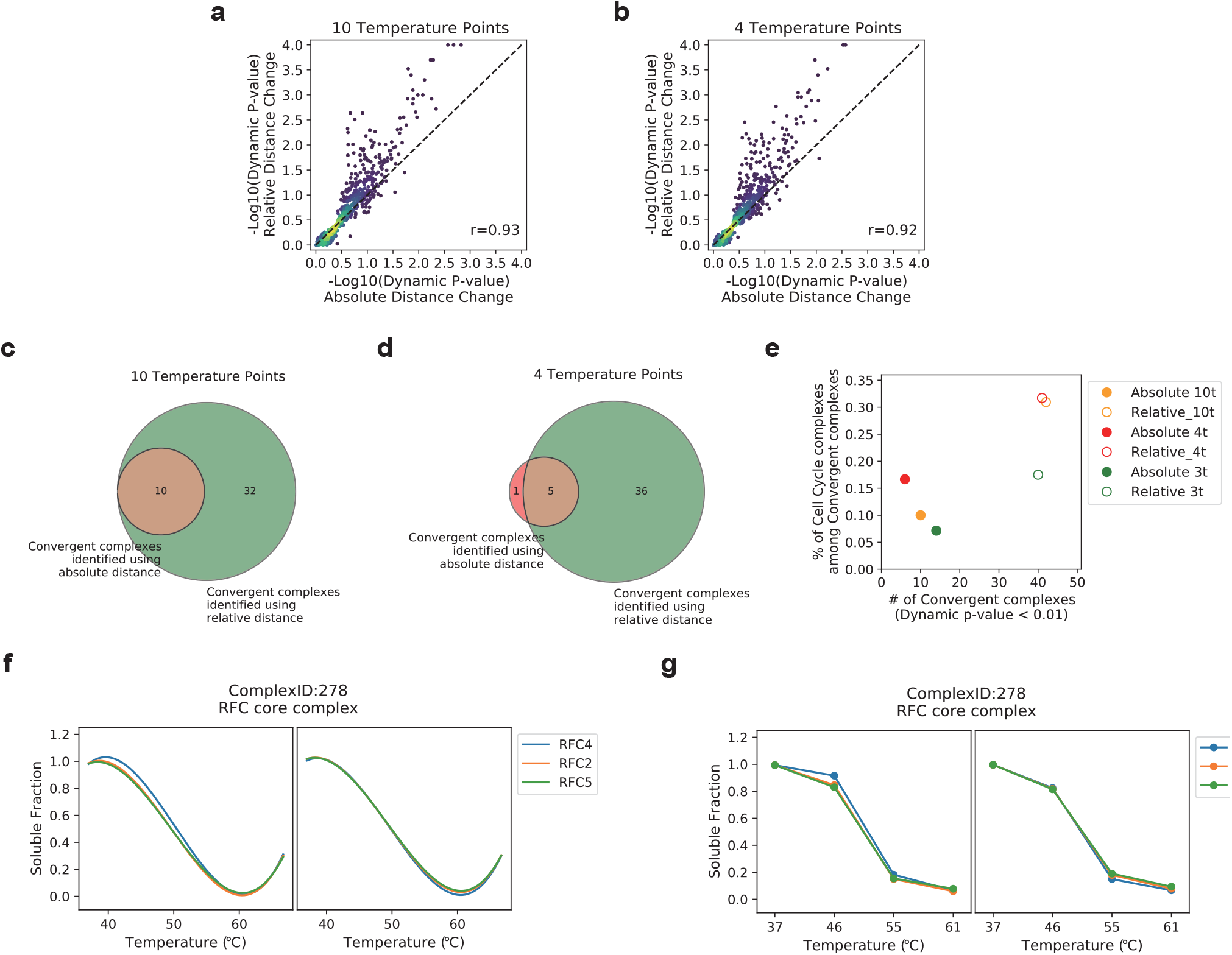
Relative distance algorithm identifies more dynamic complexes in TPCA and Slim-TPCA. **(a-b)** Correlation of the dynamic TPCA p-values of CORUM complexes obtained using the relative distance algorithm and the absolute distance algorithm for 10 and 4 temperature points, respectively. **(c-d)** Comparison of the dynamic complexes identified by the relative distance algorithm with the absolute distance algorithm at 10 and 4 temperatures, respectively. **(e)** Number and functional class of convergent protein complexes identified with different algorithms. Compared to the absolute distance algorithm, the relative distance algorithm identifies more dynamic complexes associated with cell cycle, according to the annotation in the CORUM database. **(f-g)** Curves of subunits in Complex 2792 under DMSO and MTX, with 10 temperature points and 4 temperature points, respectively.

As expected, the relative distance algorithm identified more modulated protein complexes than absolute distance algorithm across 10, 4 and 3 temperatures tested. In particular, the modulated protein complexes identified by relative distance scoring algorithm encompassed all complexes identified by absolute distance scoring algorithm with 10 temperature points (Fig.3c), and all but one complexes when using 4 temperature points (Fig.3d). Importantly, the relative distance algorithm also identified a higher percentage of protein complexes associated with cell cycle and DNA processing (according to the annotation provided by the CORUM database) which is more in accordance with the experimental condition in which samples are collected where cells are trapped in the S-phase of cell cycle (Fig.3e). For example, the RFC complex, which is involved in DNA replication, is identified to be dynamically modulated with the relative distance algorithm but is missed by absolute distance algorithm (Fig.3f-g). This suggest that the relative distance algorithm has improved sensitivity in identifying modulated protein complexes.

### Optimization of P-value algorithm using fitted Beta distribution

To quantify the significance of TPCA signature observed as well as its changes across conditions, we had adopted a bootstrapping approach to estimate p-value which is the probability observing random complexes with similar or more extreme values (either absolute TPCA signature or changes). Specifically, the bootstrapping (sampling) algorithm compares the average distance or change in average distance of the complex with 10,000 randomly sampled protein sets of the same size. This method obtains discrete p-values and takes up considerable time. Here, we optimized this part of TPCA workflow fitting a specific data distribution with smaller sample size for estimating p-value.

We validated this approach with the intact cell data of K562 published in the proof-of-concept paper with 4 temperature points of 37°C, 46°C, 55°C, 61°C recommended earlier. First, using the Chi-square statistics, we observed that Beta distribution generally fitted well to the distributions of Manhattan distance derived from bootstrapping algorithm across different number of temperatures for both absolute and relative distance scoring (Supplementary Fig.4). Next, we verified that Beta distribution fitted with 100, 200, 500, and 1,000 random complexes is similar to actual distribution of Manhattan distance computed from 10,000 random complexes, indicating that small number of samples fitted to Beta distribution can be used in place of excessive sampling. Thus, we tested the Beta distribution fitting algorithm with the dataset of K562 cells trapped in S-phase of cell cycle using MTX. We observed more dynamically modulated protein complexes being identified with this approach, that is generally accompanied with a slight increase in the proportion of protein complexes related to cell cycle and DNA processing across 10, 4 and 3 temperature points (Fig.4d).

**Fig. 4.**
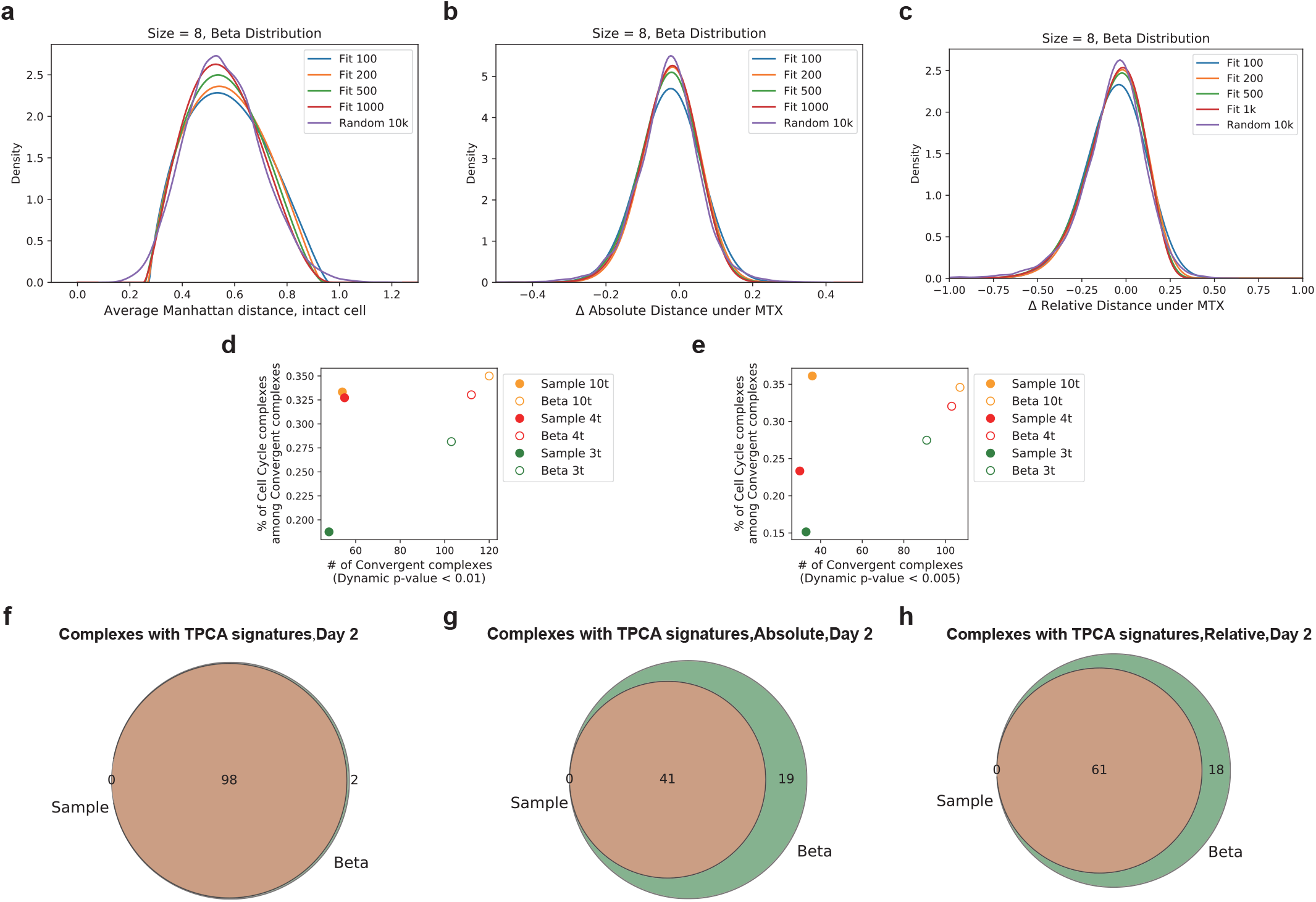
Optimization of P-value estimation algorithm using fitted Beta distribution. **(a-c)** The algorithm with small number of samples combined with the fitted Beta distribution can simulate the distribution of large number of samples very well as observed for Manhattan distance (A), Δ absolute distance (B) and Δ relative distance (C). **(d-e)** Number and functional class of convergent protein complexes identified based on new P-value estimation algorithm. Beta distribution fitting algorithm identifies more dynamically modulated complexes with higher percentage associated with cell cycle and DNA processing generally. **(f-h)** The result of complexes with TPCA signatures and dynamic modulations obtained by the new algorithm maintain a good agreement with those obtained by the traditional algorithm with massive sampling based on Manhattan distance (F), Δ absolute distance (G) and Δ relative distance (H).

Subsequently, we cross-validated the new algorithm on the viral infection dataset^1^ with 4 temperature points of 36.9°C, 46.6°C, 55.3°C, and 61.2°C. We identified the complexes with significant TPCA signatures or differentially modulated during viral infection and compared them with the complexes identified with the original sampling method. The new algorithm of fitting Beta distributions essentially recovered all the protein complexes identified by the traditional sampling algorithm (Fig.4e-g) regardless whether Manhattan distance, difference in absolute Manhattan distance or difference in relative Manhattan distance is used. Importantly, this new algorithm requires less than 1% of the time needed in the original sampling processing with 10,000 random complexes.

### Slim-TPCA identifies dynamically modulated complexes in response to glucose deprivation

Cancerous cells are often found to have altered energy metabolism where ATP are preferably generated from less efficient glycolysis in cytoplasm over oxidative phosphorylation pathway in mitochondria. Glucose, a key energy source that is supplied abundantly in most culture media, is consumed excessively, and found to induce cell death in many cancerous cells when withdrawn (glucose addiction)^34^. However, not all cancerous cells exhibit dependency on glucose for survival and proliferation. Glutamine, another key nutrient consumed excessively by cancerous cells, could be a source for ATP production through its α-ketoglutarate intermediate that feed into the tricarboxylic acid (TCA) cycle^35^. Here, we incorporate all the modifications and enhancements made in Slim-TPCA to investigate the dynamics of CORUM-annotated protein complexes under glucose deprivation which we applied on K562 suspension cells to minimize perturbation to cell morphology and cellular physiology during cell harvesting (Supplementary Fig.5-12).

First, we observed minimal apoptosis in K562 cells at different duration of glucose deprivation tested based on FACS analysis (Supplementary Fig.13). Nevertheless, replication of cells was stalled in the first 24 hours with resumption observed after 48 hour of glucose deprivation, indicating successful adaptation of K562 cells to glucose-deficient condition. Accordingly, K562 cells were harvested at 0th, 4th, 8th, 24th and 48th hour of glucose deprivation. At each time point, vials of cells were subjected to the optimal set of 37°C, 46°C and 55°C identified previously, and processed with TMT16 reagents for concurrent analysis on mass spectrometer. Three sets of biological replicates are performed, and the data collected are pooled for integrative analysis to maximize signal and reduce stochastic variation. Overall, we observed high data reproducibility with average Pearson’s correlation of 0.76 and 0.82 in protein solubility at each time point for 46°C and 55°C respectively across the three biological replicates (Supplementary Fig.14 and Supplementary Fig.15).

TPCA signature permits the identification of both increasingly associated (convergent) and increasingly dissociated (divergent) protein complexes in situ compared to a baseline. Here, glucose-deprived K562 cells are compared to cells at on start of glucose deprivation (0th hr). On average, we observed about twice more divergent than convergent protein complexes across all the time points profiled (Fig.5b), arguably in accordant with the downregulation expected for most basal cellular processes. Indeed, we observed proteins in divergent protein complexes over-represented in translation and ribosome biogenesis from samples at 4th, 8th, 24th but not at 48th hour of glucose deprivation (Fig.5c and Supplementary Fig.16).

**Fig. 5.**
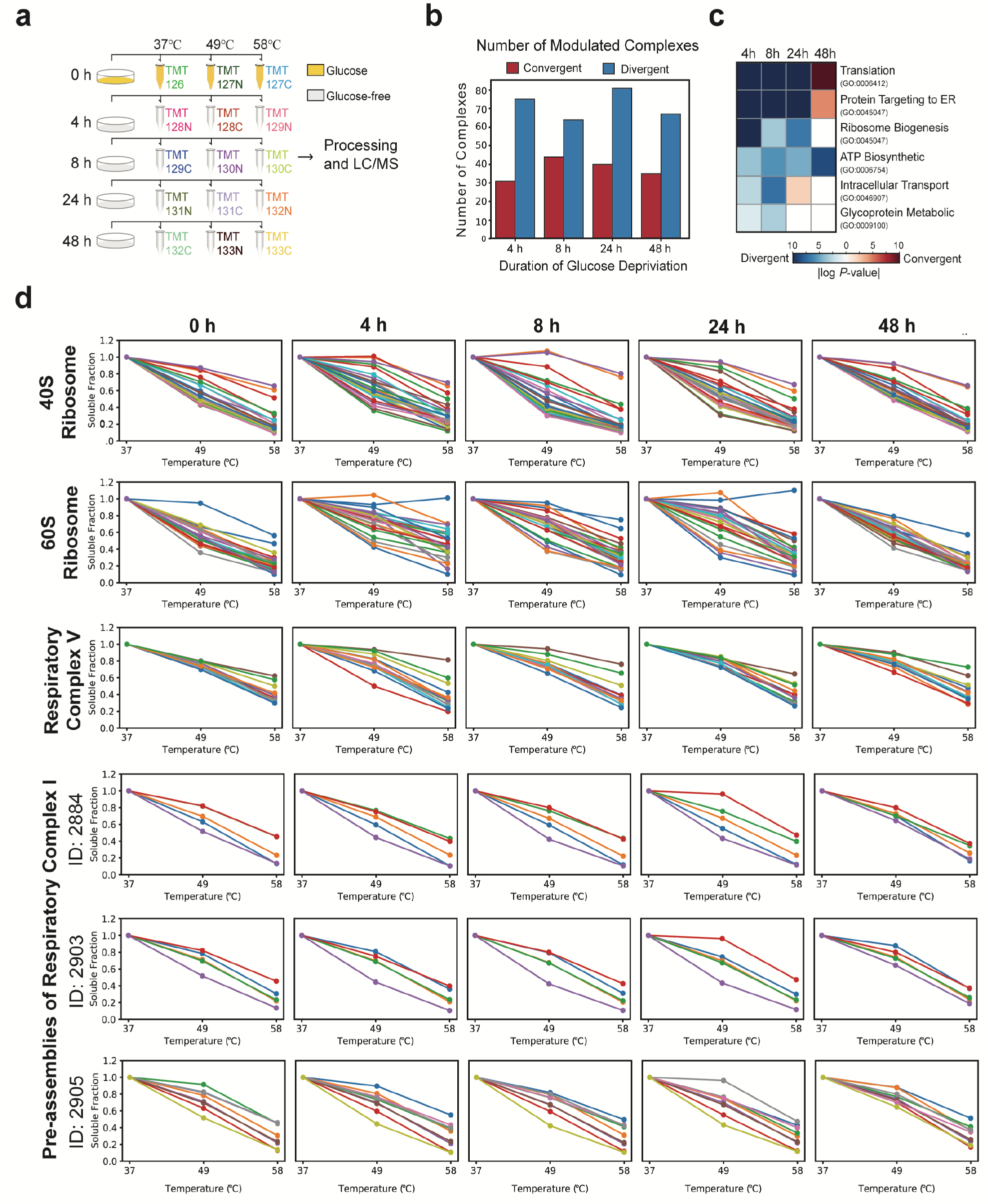
Application of optimized TPCA profiling to K562 cells under glucose deprivation. **(a)** Schema of samples collected. K562 are deprived of glucose in culture medium and harvested at different time point for heating at 3 temperatures for TPCA profiling. **(b)** Numbers of CORUM-annotated protein complexes detected at different time point with convergent and divergent TPCA signature. **(c)** Major functional classes of proteins in divergent protein complexes. **(d)** TPCA profile of selected convergent and divergent protein complexes at different time point of glucose deprivation.

Thus, we inspected the previous observation more closely with the TPCA profile of 40S and 60S ribosomes at different time point of glucose deprivation. The thermal solubilities of both 40S and 60S ribosomal proteins are most dissimilar to each other (indicating dissociation of complexes based on TPCA theory) at 4th, 8th, 24th hour of glucose deprivation. However, the TPCA profile of ribosomal proteins at 48^th^ hours of glucose deprivation are highly similar to data collected from samples without glucose deprivation suggesting resumption of translational activities. Previous studies had revealed system-wide attenuation in protein translation under glucose deprivation^36,37^ that occurred independent of mRNA abundance^38^. Thus, the disassembly of ribosomes could possibly contribute or arise from decreased protein translation, an energy-intensive process^39,40^ which prerequisite in many cells is the availability of glucose^37^. Interestingly, we also observed similar pattern of divergent and convergent TPCA signatures for the 28S and 39S ribosomal subcomplexes localized in mitochondria (Fig.6a) suggesting cell-wide attenuation of protein translation. This demonstrated the utility of new streamlined TPCA protocol for identifying modulated protein complexes including those localized in membrane-bound organelles.

**Fig. 6.**
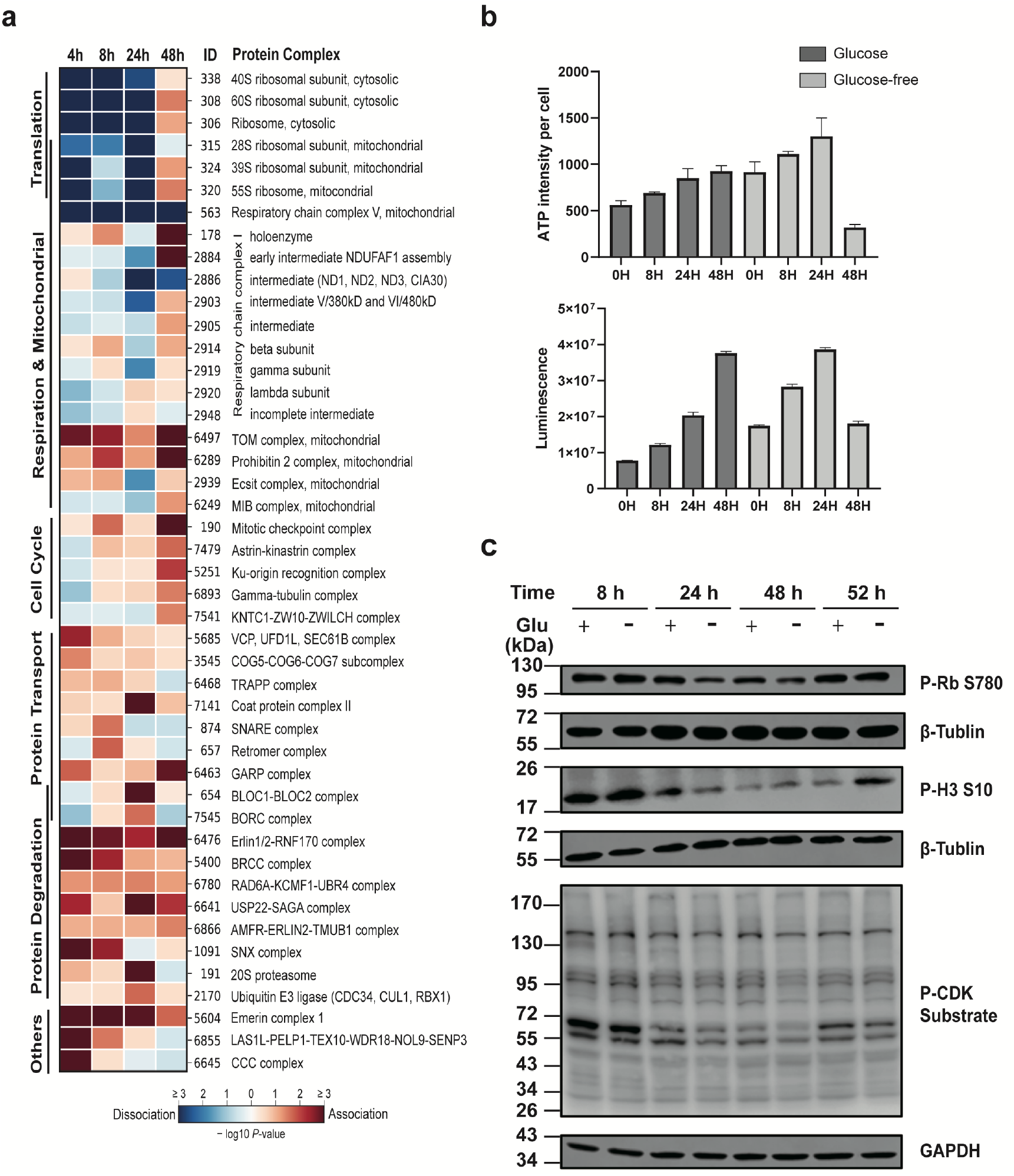
Glucose deprivation induces protein complexes changes in K562 cells. **(a)** Dynamics of CORUM-annotated protein complexes under glucose deprivation as profiled by Slim-TPCA. Complexes are grouped based on their functions. **(b)** K562 cells were incubated in presence and absence of glucose followed by quantification of ATP based on luminescence intensity (n = 3 independent biological experiments). Data are presented as mean values ± s.d. (error bars). **(c)** K562 cells were incubated in presence and absence of glucose, harvested at different time points and analyzed by immunoblotting using Phospho-Rb (Ser780) antibody, Phospho-Histone H3(Ser10) antibody, and Phospho-CDK Substrate antibody. GAPDH antibody and β-Tubulin antibody were used as loading control.

Intuitively, the deprivation of glucose could potentially alter mitochondrial protein complexes involved in cellular respiration. Thus, we focus on validating our new streamlined TPCA protocol on such protein complexes annotated in the CORUM database. On average, less than 20% of core set of CORUM-curated protein complexes are found modulated at any time point profiled. Encouragingly, most mitochondrial protein complexes involved in respiration (consisting of respiratory chain complex V, the various subcomplexes and intermediates of respiratory chain complex I) exhibit differential TPCA signature under glucose deprivation.

In particular, the respiratory chain complex V (annotated as F1F0 ATPase complex in CORUM database) exhibits very consistent divergent TPCA signature that is strongest in the first 8 hours but weaken thereafter. Based on TPCA profile, there seem to be a recovery of the complex sometime after 24 hour of glucose deprivation (Fig.5d). On other hand, the various intermediates and subcomplexes of respiratory chain complex I exhibit more diverse and muted but nevertheless statistically significant changes in TPCA signatures. Interestingly, while most of these intermediates and subcomplexes manifest divergent TPCA signatures that peaked at 24^th^ hour, the TPCA signature were reversed (convergent) at 48^th^ hour.

Similarly, divergent TPCA signature was observed for the MIB (mitochondrial intermembrane space bridging) complex (Fig.6a) formed between the MICOS (mitochondrial contact site) complex and the SAM (sorting and assembly machinery) complex located in the inner and outer membrane respectively^41,42^. These complexes are involved in the formation of cristae and junction that optimize the microenvironment for assembly of respiratory chain complexes and ATP production. Like the various intermediates and subcomplexes of respiratory chain complex I, the TPCA signature of the MIB complex also diverged in the first 24 hours of glucose deprivation but subsequently converged based on samples collected at 48^th^ hour of glucose deprivation.

Overall, TPCA profiling suggests that cellular respiration is muted in the first 24 hours with sign of recovery thereafter based on samples collected at 48^th^ hour of glucose deprivation. To investigate this further, we quantified the amount of ATP in K562 cells at different time point of glucose deprivation. Somewhat unexpected initially, we observe small but gradual increase in total ATP and average ATP abundance (per cell) in the first 24 hours of glucose deprivation with a sharp decrease in ATP in cells collected at 48^th^ hour of glucose deprivation (Fig.6b). Thus, ATP abundance seems to correlate inversely with TPCA signatures of the various respiratory complexes. We reasoned this could arise from the huge ATP demand of cells in many basal cellular activities after successful adaptation to glucose-deficient condition. Indeed, this seems to be supported by the observed convergent of 40S and 60S ribosomal protein complexes at 48^th^ hour of glucose deprivation suggesting increased protein translation which arguably consume a large majority, if not most, of cellular ATP^39,43^. Overall, we observed that when ribosomal assembly are presumably downregulated (based on TPCA signature), higher cellular ATP abundance was observed.

As most of the identified modulated protein complexes dissociated and are involved in basal cellular processes that we will reasonably expect in cells deprived of a source of energy, we focus on analyzing convergent protein complexes to identify those potentially implicated in metabolic reprogramming of cells in the absence of glucose. While the various mitochondrial respiratory complexes mostly exhibit divergent TPCA signatures, other differentially modulated mitochondrial protein complexes manifest a strong convergent TPCA signature instead. They are the TOM complex involved in mitochondrial protein import, the Prohibitin 2 complex that inhibit caspase-dependent apoptosis^44^, and the ECSIT (evolutionarily conserved signaling intermediate in Toll pathways) complex implicated in the regulation of mitochondrial respiration and mitophagy^45^ (Fig.6a).

In addition, we observed increased association of many protein complexes involved in protein transport and vesicle formation with peaked convergent TPCA signatures observed at 8th and 24th hour time point sampled that included BLOC and BORC protein complexes associated with lysosome. At the same time, we observed convergent protein complexes implicated in degradation of various proteins throughout the time points profiled. These complexes include the Erlin1/2-RNF170 complex and the SNX complex implicated in degradation of IP3 (Inositol 1,4,5-trisphosphate) receptors and EGFR (epidermal growth factor receptor) respectively, as well as various protein complexes that possesses E3 ligases (e.g. the BRCC complex and the AMFR-ERLIN2-TMUB1 complex). Correspondingly, we observed convergence of 20S proteasome and an E3 ligase complex (consisting of CDC34, CUL1 and RBX1) that peaked at 24^th^ hour of glucose deprivation. Hence, there seems to be an increase in protein degradation activities perhaps to free amino acids for energy metabolism.

We also observed a few protein complexes that converge only at 48^th^ hour and are implicated in chromosome segregation and mitosis. These complexes include both the mitotic checkpoint complex and the astrin-kinastrin complex that are involved in correct alignment of chromosome during mitosis, the gamma-tubulin complex (CORUM ID: 6893) that associates with centrosome involved chromosome segregation, and the KNTC1-ZW10-ZWILCH complex that is associated with kinetochores on chromatids. Collectively, the convergent of these complexes suggest possible onset of mitosis around 48^th^ hour of glucose deprivation. As such, we performed immunoblotting for specific mitotic protein marker namely phosphorylation of histone H3 and retinoblastoma protein at S10 and S780 respectively. We observed decrease of these markers at 24th and 48^th^ hour but an increase at 52^nd^ hour of glucose deprivation, suggesting the protein complexes implicated in chromosome alignment and segregation converge before activation of mitosis-associated signaling. We also compared the phosphorylation levels of CDK substrates, another mitotic protein marker, by immunoblotting. Similar to what was observed for phosphorylation of histone H3 and retinoblastoma protein, we observed reduced CDK substrate phosphorylation before 48^th^ hour of glucose deprivation that increased at the 52^nd^ hour indicating possible onset of mitosis (Fig.6c).

## Discussion

Protein complexes are the key effectors in most cellular activity that need to be dynamically assembled and dissolved according to cellular needs. Unlike most proteome-wide experimental methods, the intracellular assembly state of protein complexes can be profiled in situ with TPCA rapidly. The method had been verified in primary cell and tissues and had been deployed to identify protein complexes modulated during cell cycle, viral infection and T-cell infection. Conceptually, the method could be readily deployed across various cellular conditions to profile the assembly state of uncharacterized protein complexes identified from large scale projects to aid in their functional annotation.

Here, we introduce Slim-TPCA, an improved TPCA method using fewer temperature points and improved data deconvolution algorithms offering a series of improvements in terms of experimental design and data analysis over previous method. We showed that less temperature points can be used with minimal or negligible lost in TPCA signal. This permits more flexibility in the experimental design, allowing more time points or experimental conditions per analysis while reducing sample needed, experimental cost and batch effect in mass spectrometry analysis. At the same time, the new data processing algorithms offers over 100X reduction in time required for statistical calculations while identified modulated protein complexes with improved sensitivity.

In this work, we had identified the optimal smaller sets of temperatures for TPCA profiling of mammalian cells. The temperature points identified fitted well with the expectation that reasonably spaced temperatures maximally captured TPCA signature. For the application of TPCA profiling in other species, similar principle could be applied for selection of optimal set of temperatures. We had also adopted Manhattan distance for encapsulating TPCA signal instead of Euclidean distance used in the original TPCA work as Manhattan distance gave the best overall performance across all the number of temperatures tested. Although Pearson’s correlation performs as well as Manhattan distance, its ability to encapsulate TPCA signature dimmish more drastically with less data points compared to Manhattan distance and other metrics. Another major deviation from the previous TPCA format is the use of relative change in TPCA signature over absolute difference to better characterize the dynamics of protein complexes, especially those with already strong inherent or basal TPCA signatures.

Conceptually, the use of relative distance to quantify changes in TPCA signature across conditions can reduce false negative to identify more modulated protein complexes but the approach can also lead to a higher false positive rate. The latter is particularly more pronounced if quality of the data is noisier. We previously reported that precision or reproducibility in thermal solubility quantified depends in part on number of peptide-spectral match (PSM) and ion intensity of proteins^14^. To reduce false positive rate, proteins will less precise values can be filtered based on these criteria. Alternatively, data from multiple technical and biological replicates can be integrated to obtain averages that are nearer to the true values. We prefer the latter approach as the former approach of filtering based on PSM and ion intensity can remove many proteins. Using data integrated from three biological replicates reported in previous work, we found the relative distance identified more modulated protein complexes that correlate with expected biological activities.

Another improvement made is over 100X reduction in time needed to compute the statistical significance of TPCA signature and its changes across conditions. Here, we adopted the strategy of using lesser sampling (500 rather than 10,000 sets of randomly selected protein) fitted to Beta distribution with computed p-value that correlate well with empirical p-value. This improvement is provided as an option in our provided Python package and ProSAP GUI software.

Finally, we combined all the improvements and changes made applying Slim-TPCA to profile K562 cells at five time points of glucose deprivation using three temperature points which are performed in a set of TMT experiment for each replicate. Encouragingly, we identified modulated protein complexes involving in biological processes that are either reportedly or expectedly associated with glucose starvation. Our profiling identified increased dissociation of both cytoplasmic and mitochondrial ribosomes that is coherent with reported attenuation of protein translation during glucose starvation. Many mitochondrial protein complexes involved in or associated with respiration and complexes implicated in vesicle-based protein transport are found to be modulated revealing that TPCA is applicable to protein complexes localized within membrane-bound organelle and those associated with membrane. In summary, we envision Slim-TPCA and associated software package to expediate functional characterization of existing and newly identified protein complexes.

## Materials and Methods

### Cell culture

K562 cells were cultured either in RPMI 1640 medium (Gibco) or glucose-free RPMI 1640 medium (Gibco) supplemented with 10% FBS (PAN) and 1% penicillin**−**streptomycin (Gibco) at 37 °C, and 5% CO2 in a humidified environment. Cells were washed twice with ice-cold PBS and subsequently subjected to (1X) lysis buffer containing a final concentration of 50 mM HEPES pH 7.5, 10 mM MgCl2, 5 mM β-Glycerophosphate (sodium salt hydrate), 1 mM Tris(2-carboxyethyl) phosphine hydrochloride (TCEP), 0.1 mM Sodium orthovanadate,0.2%(w/v) n-dodecyl β-D-maltoside (DDM), and EDTA-free protease inhibitor ((Sigma-Aldrich, USA). Cell suspension was subjected to five times flash-freezing in liquid nitrogen and rapid thawing in water to facilitate cell lysis. After centrifugation at 21000g for 20 min at 4 °C, the supernatant was transferred to a new tube and the protein concentration was measured by the BCA assay kit (Thermo Fisher Scientific, USA).

### Preparation of MS samples

Samples were prepared by SISPROT^46,47^, a spintip-based device, as previously described. Briefly, SISPROT device was fabricated by packing several plugs of C18 disk (3 M Empore, USA) into a standard 200 μL pipette tip and then introducing certain amounts of mixed beads. The mixed beads are composed of POROS SCX beads and POROS SAX beads (Applied Biosystems, USA) in a ratio of 1:1. Subsequent steps of sample loading, protein reduction, alkylation, digestion, TMT pro ple(Thermo Fisher Scientific, USA)-labeling and desalting were all done on the SISPROT tip. The TMT mixed sample was dried using a vacuum centrifugal evaporator and redissolved with 1% (v/v) formic acid followed by fractionating into six fractions with a stepwise increasing gradient of ACN. Finally, eluted peptide samples were lyophilized to dryness and redissolved in 0.1% (v/v) formic acid in water for nano-LC-MS/MS analysis.

### MS Analysis and Data Analysis

Each sample was diluted with 0.1% FA prior to separation on 20 cm x 100 um EASY-Spray C18 LC column with a 135 min gradient on an UltiMate 3000 HPLC system (Thermo Fisher Scientific). The mobile phase was solvent A (0.5% acetic acid in water) and solvent B (80% ACN, 0.5 % acetic acid in water). Data were acquired by Orbitrap Exploris 480 mass spectrometer (Thermo Fisher Scientific): MS1 scan resolution was 60,000 and MS/MS scan resolution was 30000 using turbo TMT. Raw files were searched using Proteome Discoverer (PD) software (Version 2.4, Thermo Fisher Scientific) against the human proteome fasta database (Uniprot,20376 entries, downloaded on May 03, 2022). The maximum missed cleavage for trypsin digestion was set to 2. The mass tolerance for peptide precursors was 10 ppm and the mass tolerance for fragment ions was 0.02 Da. Carbamidomethyl of cysteine residues, TMT pro modification of lysine residues and TMT pro modification of peptide N-terminal were selected as fixed modifications while oxidation (M) and deamidation (NQ) were selected as variable modifications. FDR control is performed at 0.01 level at the PSM and peptide levels.

### Flow cytometry detection of apoptosis

Cell apoptosis was detected using Annexin V-FITC Apoptosis Detection Kit (Beyotime, China). K562 cells were harvested and washed with ice-cold PBS twice prior resuspended in 1X annexin-binding buffer at 5 million cells per tube. Cells were then stained with FITC Annexin V and PI solution protecting from light at room temperature for 15 minutes and analyzed by flow cytometry (BD FACSCanto SORP) as soon as possible.

### Detection of ATP levels

ATP measurement was determined by CellTiter-Glo Luminescent Cell Viability Assay (Promega, USA). K562 cells under glucose or glucose-free conditions for 0 h,8 h, 24 h, and 48 h were seeded in a 96-well plate at 10 thousand cells per well. Cells were lysed using Cell Titer-Glo® reagent and mixed for 2 min on an orbital shaker. The plate was incubated for 10 min and analyzed by microplate reader (EnSpire), while cells number were measured by blood counting plates.

### Treatment with Pharmacological Inhibitors

K562 cells under glucose or glucose-free conditions for 8 h, 24 h, 48 h, and 52 h with and without Chloroquine (10 uM) for 4 h were harvested. Cells were lysed using RIPA buffer containing a final concentration of 50 mM Tris-HCL pH 8.0, 150 mM NaCl, 1% Triton x-100, protease cocktail, and 1 mM PMSF facilitated by freeze-thawing five times using liquid nitrogen. After centrifugation at 21000g for 20 min at 4 °C, the supernatant was transferred to a new tube and the protein concentration was measured by the BCA assay. Cell lysates were analyzed by western blotting.

### Antibodies

Phospho-Rb (Ser780) antibody, Phospho-Histone H3(Ser10) antibody, Phospho-CDK Substrate antibody, and β-Tubulin antibody were obtained from Cell Signaling Technology (Danvers, USA). GAPDH antibody was purchased from Proteintech (Rosemount, IL, USA). HRP-labeled Goat Anti-Rabbit IgG(H+L) was obtained from Beyotime (Nanjing, China).

### Western blotting

Twenty micrograms of total protein from each sample were resolved on SDS-polyacrylamide gels and transferred to polyvinylidene fluoride (PVDF) membranes. After blocking in phosphate-buffered saline containing 2.5‰ Tween 20 (PBST) and 5% BSA for 1.5 h, the membranes were incubated with the indicated primary antibodies overnight at 4 °C. Then, the membranes were incubated with HRP-labeled Goat Anti-Rabbit IgG(H+L) for 1 h at room temperature. The relative density of each band was analyzed on an Odyssey infrared scanner (LICOR Bioscience, Lincoln, NE, USA). GAPDH and β-Tubulin were used as loading control.

### MS Data normalization

The thermal solubility of protein was obtained by dividing its abundance at different temperatures by its abundance at 37°C. These readings of every protein from each MS run is derived from the average reading of their isoforms (e.g. PXXXXX-1, PXXXXX-2) weighted according to the number of quantifying PSM (i.e. The number of PSMs used for quantification). To obtain more robust results, we selected proteins with 2 or more PSMs for subsequent analysis. In Slim-TPCA, because the data points were not sufficient to fit the logistic curve, we use median normalization to align data. For samples at the same temperature in different treatments/conditions, we assumed that the overall thermal stability of the proteome would not change. Let *M*_*x,t*_ *= (m*_*x,1,t*_, … *m*_*x,n,t*_) be the subset of soluble fraction for *n* proteins under treatment *x* at the temperature point *t*. For subsets *M*_*x,t*_ with the same temperature point *t* and different treatments *x*, the median *x* of medians of *n* proteins of each subset is computed. Subsequently, thermal solubility of all proteins in each subset *M*_*x,t*_ are added/subtracted with a fixed value so that all subset have the same median *x*. When using 3 temperatures, we recommend heating the samples at 37°C, 49°C and 58°C. When using 4 temperatures, we recommend heating the samples at 37°C, 46°C, 55°C and 61°C.

### Empirical statistical assessment of TPCA signature

In the conventional TPCA method, an empirical statistical assessment is used to identify complexes with non-random TPCA signatures. In Slim-TPCA, in order to identify which complexes exhibit non-random TPCA behavior, we continue the approach adopted in the original TPCA work with a few refinements. For each protein complex, we compute the average Manhattan distance (denote as *D*_*avg*_) between melting curves among all pairs of subunits of a protein complex. Then, we construct normal distributions associated with the size of the complexes, based on the mean and standard deviation of overall pairwise distance. Using these normal distributions, we transform the average distances of the complexes under test into z-scores and p-values, and use them to determine whether the complexes conform to the TPCA signature.

Specifically, let *P*_*A*_*= {p*_*1*_, *p*_*2*_, … *p*_*n*_*}* be the set of *n* unique subunits of protein complex A with solubility data. The average Manhattan distance among all unique subunit pairs (*M*_*avg*_) from *P*_*A*_is computed as

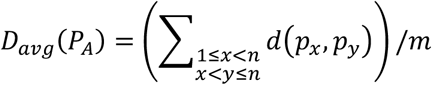

in which *m*, the number of unique subunit pairs from *P*_*A*_, is equal to (*n*^2^ **−** *n*)/2, and *d(p*_*x*_, *p*_*y*_), the Manhattan distance between melting curves of _protein_ *p*_*x*_and *p*_*y*_, is computed as

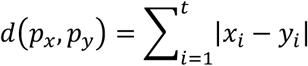

where *(x*_*1*_, *x*_*2*_, *x*_*3*_) and *(y*_*1*_, *y*_*2*_, *y*_*3*_) denote the normalized solubility of protein *p*_*x*_and *p*_*y*_at *t* temperature points.

### Empirical statistical assessment of TPCA Modulation Signature

Here, we have optimized the algorithm for identifying the dynamic modulation by calculating the changes in distance. If *P*_*A*_*= {p*_*1*_, *p*_*2*_, *…, p*_*n*_*}* is the entire set of subunits from complex A with solubility data from both Control and Treatment samples, the absolute and relative difference in distance are computed as

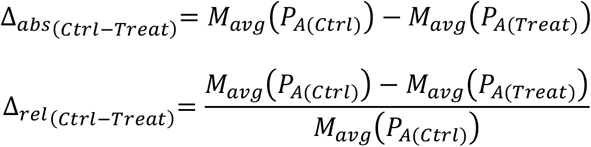

where *M*_*avg*(_*P*_*A*(*Ctrl*))_and *M*_*avg*(_*P*_*A*(*Ctrl*))_ is computed from solubility data derived from Control and Treatment samples, respectively.

To assess the statistical significance of observed Δ_(*Ctrl***−***Treat*)_, we compute Δ for 10,000 randomly selected protein set of size *n*. The TPCA Modulation Signature (P-value) of observed Δ_(*Ctrl***−***Treat*)_is then estimated as frequency of random protein set that result in Δ < Δ_(*Ctrl***−***Treat*)_. The complexes with TPCA Modulation Signature P-values less than 0.05 are identified as convergent complexes, and those with TPCA Modulation Signature P-values greater than 0.95 as divergent complexes.

### Beta distribution fitting algorithm

In empirical statistical assessment of TPCA signature and TPCA modulation signature, 10,000 random complexes of size *n* are sampled and their average distances or changes in average distances are calculated. To save computational time for high throughput data, here we use a smaller sampling size in conjunction with Beta distribution fitting instead of an excessive sampling.

We choose to sample 500 random complexes, calculate their average distance or change in average distance, and then fit them with the Beta distribution as

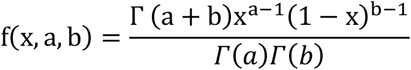

where *Γ* is the gamma function. The TPCA signature (p-value) and TPCA modulation signature (dynamic p-value) are defined as the probability of having a more extreme value in the probability density function of the fitted Beta distribution than the value of the putative complex.

## Code and Data availability

The installation package of Slim-TPCA can be downloaded directly from PyPI (https://pypi.org/project/Slim-TPCA/0.0.2/). Package documentation can be acquired online via https://slim-tpca.readthedocs.io/en/latest/index.html. If you want to test the code, testing data can be downloaded from GitHub(https://github.com/wangjun258/Slim_TPCA_examples). All the raw MS data have been deposited to the ProteomeXchange Consortium via the iProX partner repository^48,49^ with the dataset identifier PXD040078.

## Author contributions

C.T. conceived and led the project. S.S. analyzed data and wrote the algorithms. Z.Z. and F.L performed the biological experiments. J.W. and A.H. inspect and organized the code. R.T. guided the TMT-based proteomics experiment. S.S., Z.Z. and C.T. wrote the paper.

