## Supplemental Figure 1-16 for "Improved *in situ* Characterization of Proteome-wide Protein Complex Dynamics with Thermal Proximity Co-Aggregation": Suppl_Fig1-4_Slim_TPCA v20230120.docx

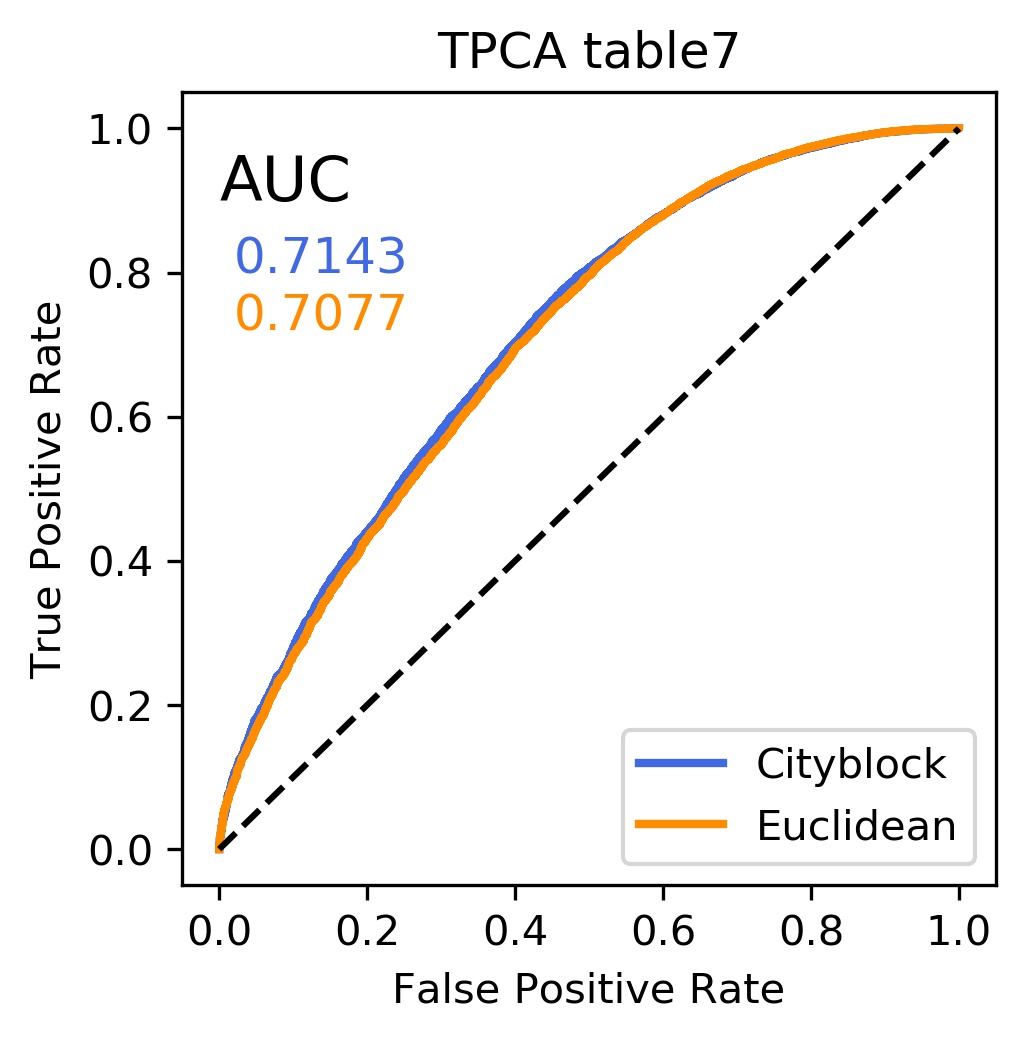

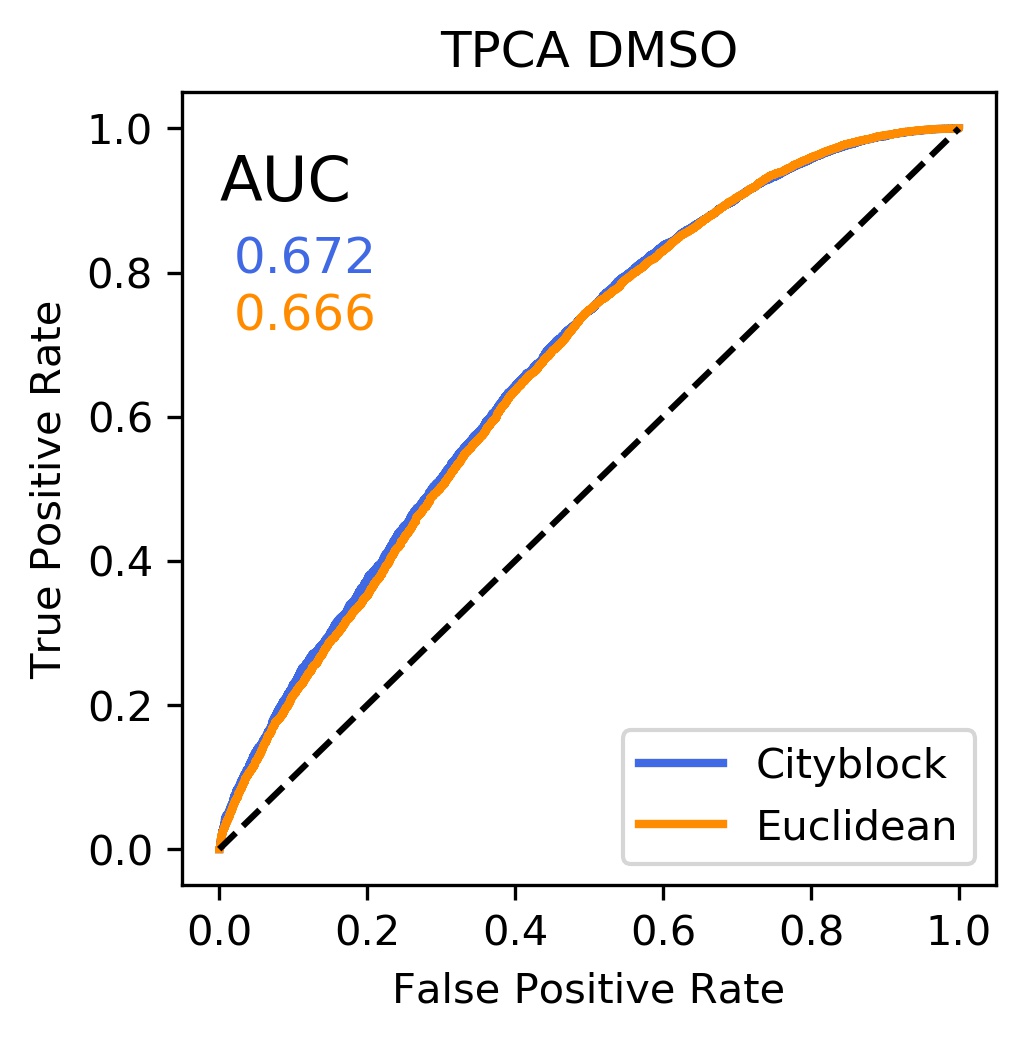

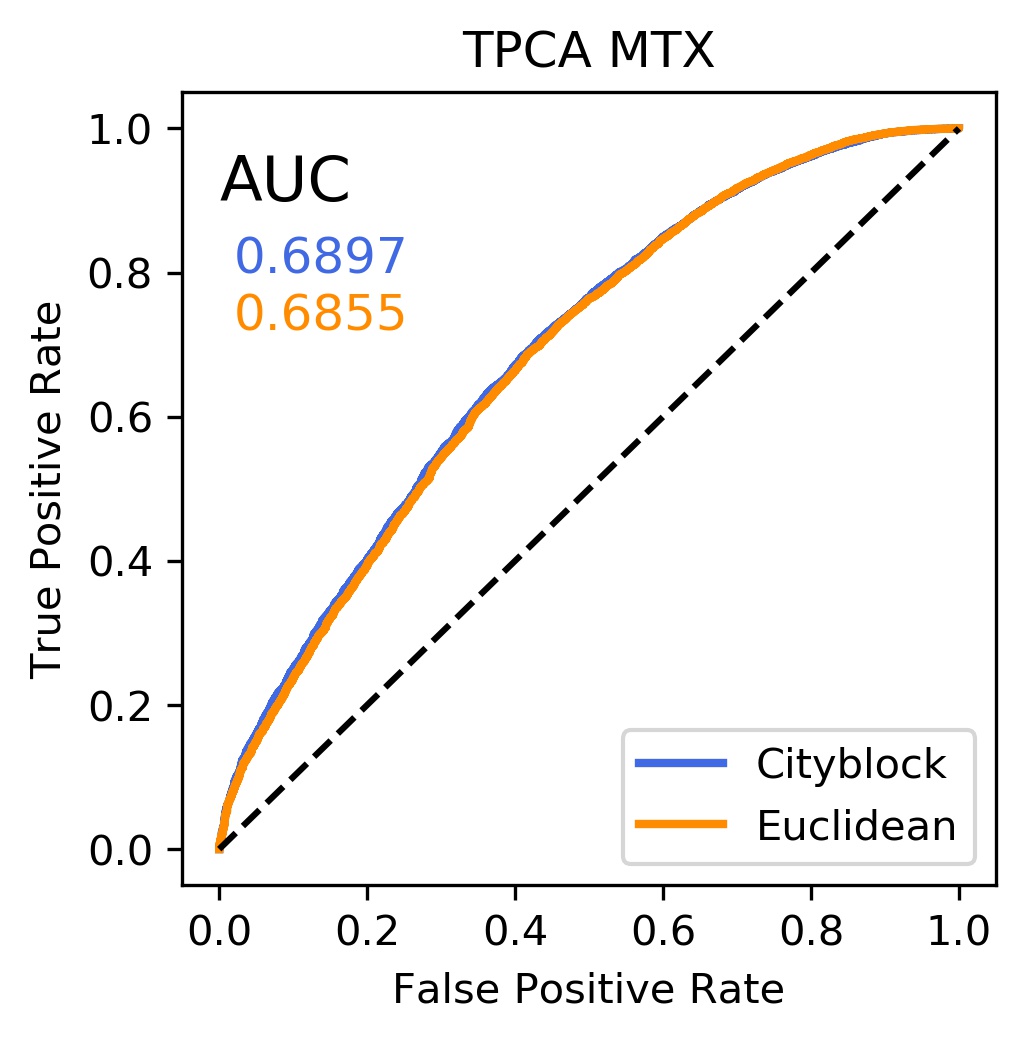

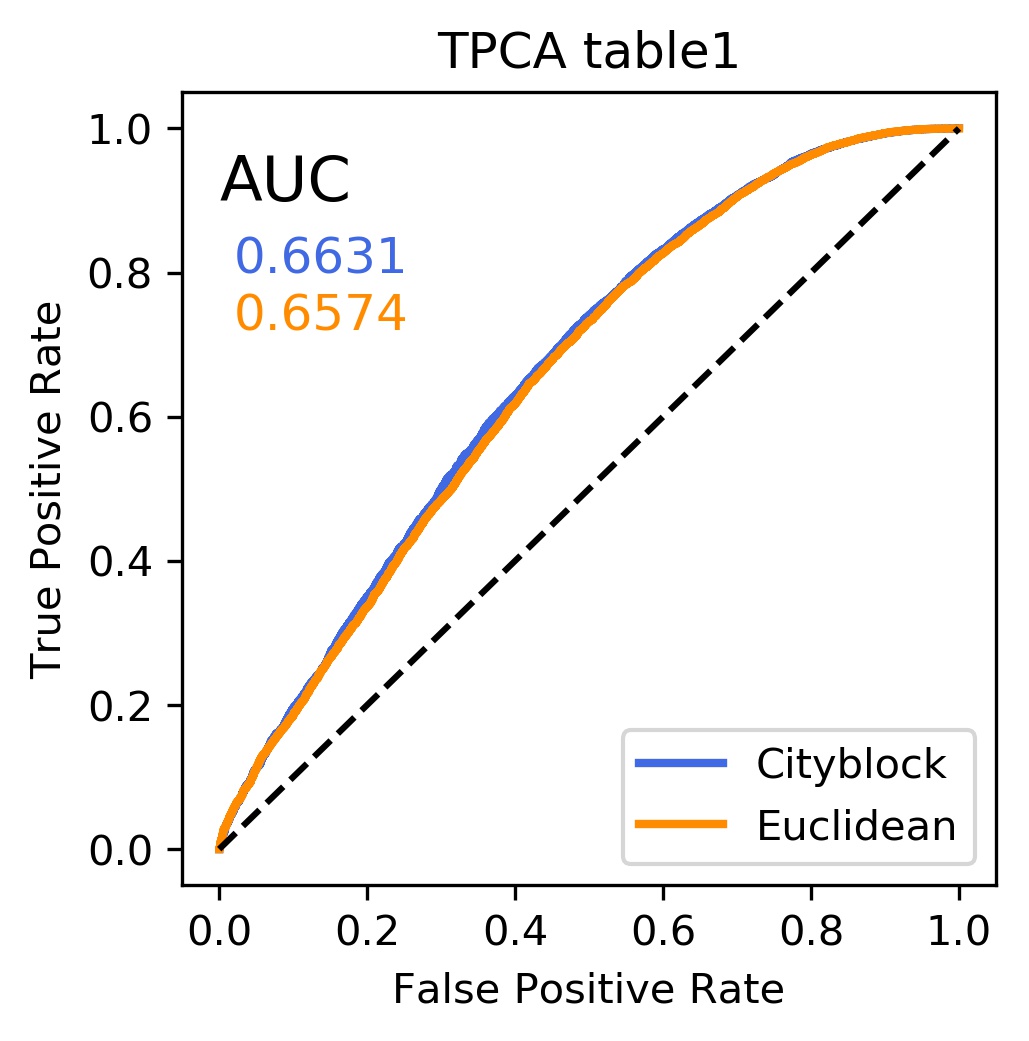

**Supplementary Fig. S1**

AUC values are improved using Manhattan distance over Euclidean distance across different data sets.

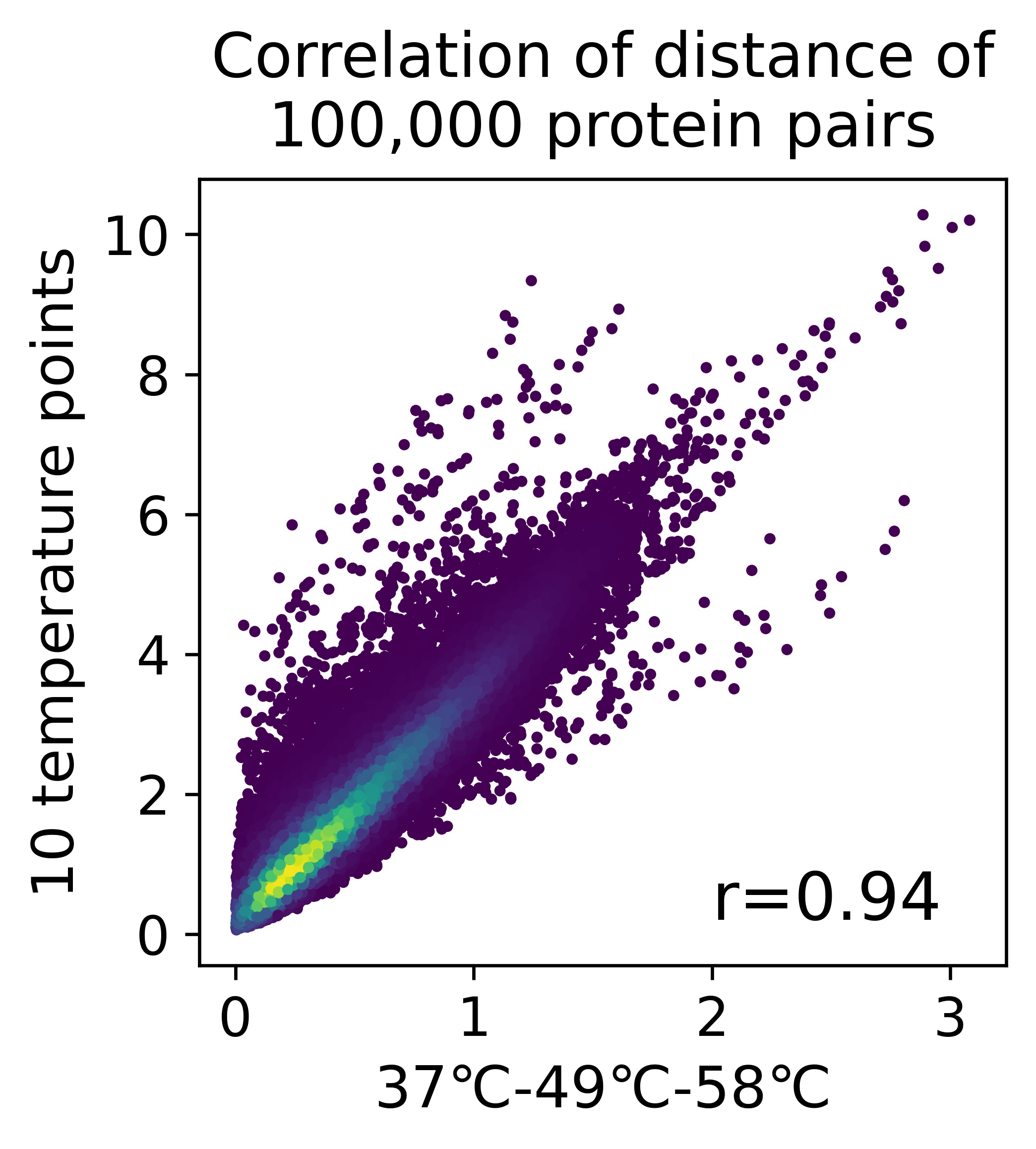

**Supplementary Fig. S2**

The protein pair distances calculated using three temperature points correlate well with those calculated using 10 temperature points.

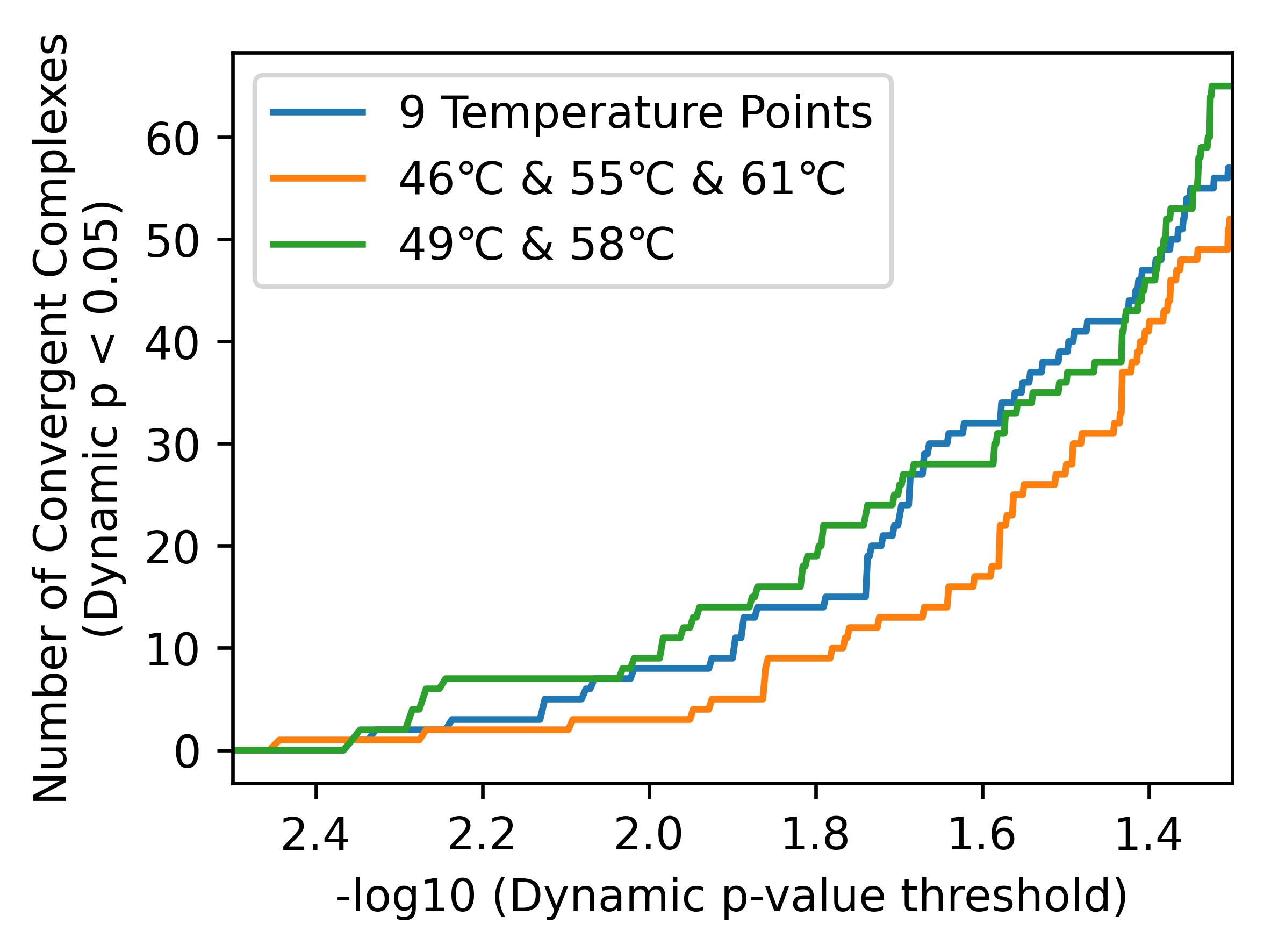

**Supplementary Fig. S3**

Dynamic complexes were identified using fewer temperature points (TPCA modulation p < 0.05).

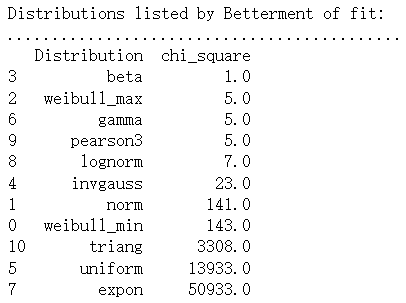

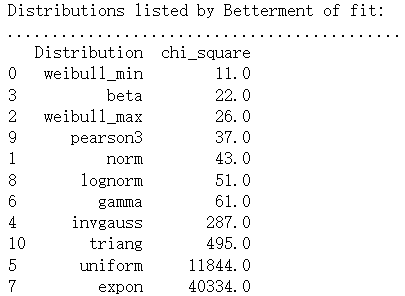

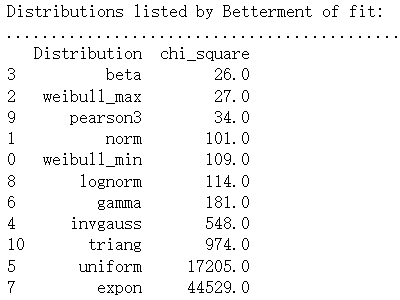

**Supplementary Fig. S4**

Beta distributions performed among the top in fitting the average distance, change in absolute distance, and change in relative distance for 100,000 complexes using data from 500 complexes, respectively.
