## Supplementary figures and images for "Improved *in situ* Characterization of Proteome-wide Protein Complex Dynamics with Thermal Proximity Co-Aggregation"

### Fig.S5_4h_convergent.pdf

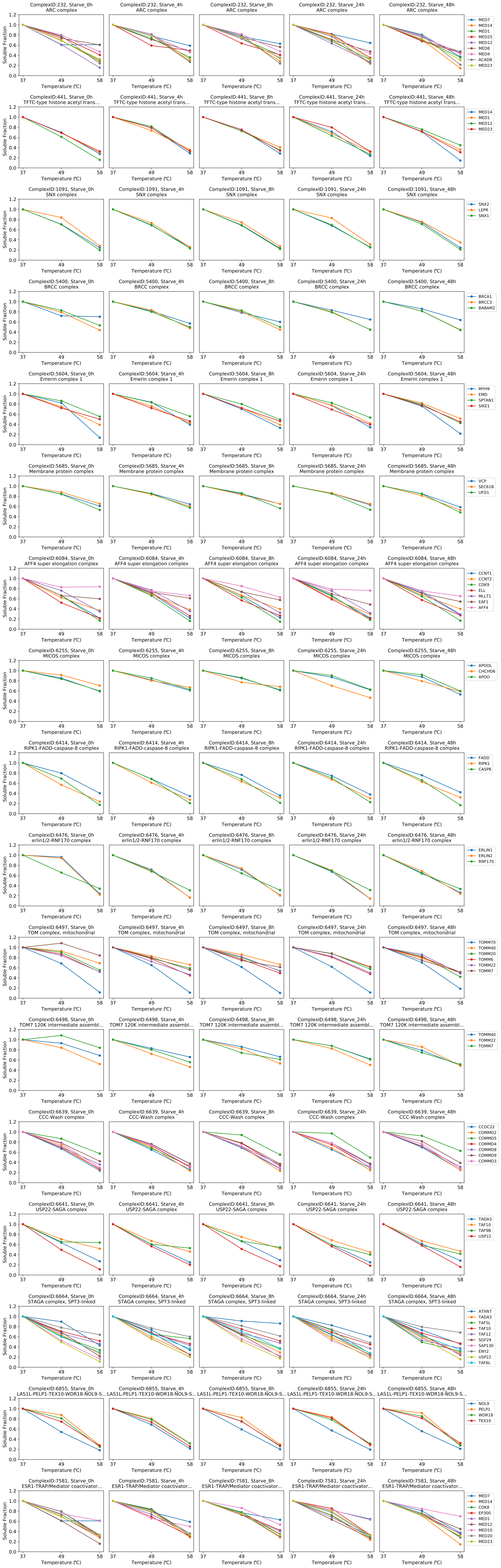

### Fig.S6_4h_divergent.pdf

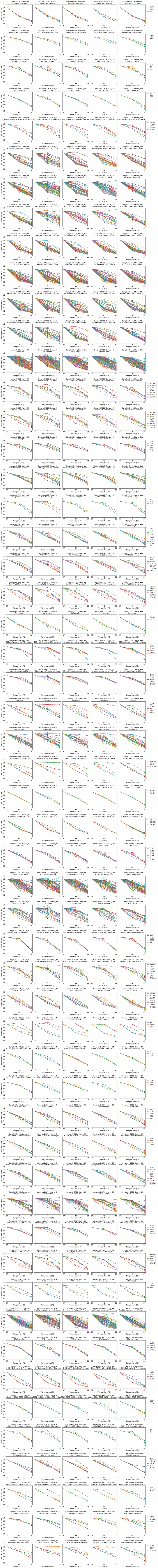

### Fig.S7_8h_convergent.pdf

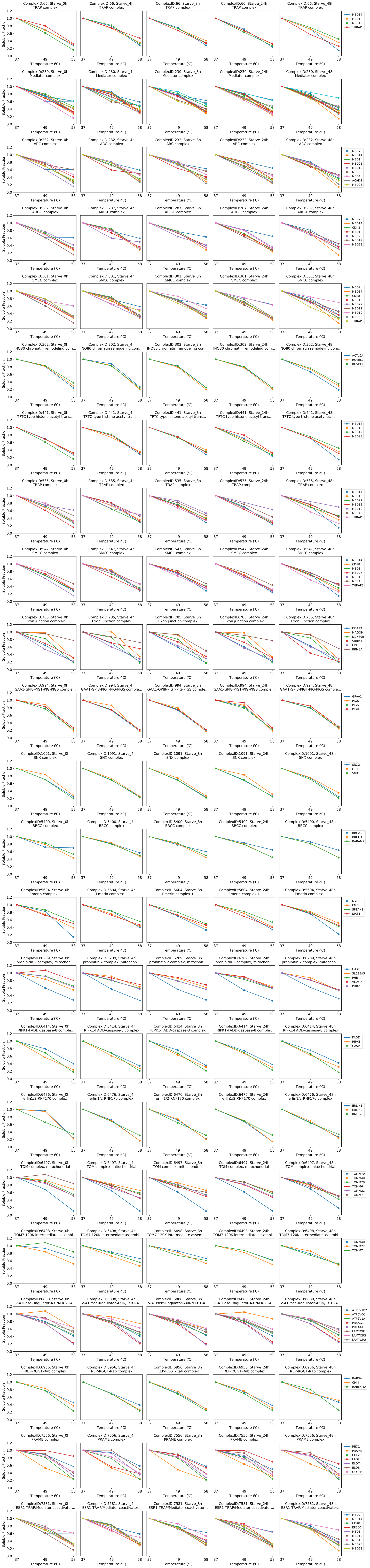

### Fig.S8_8h_divergent.pdf

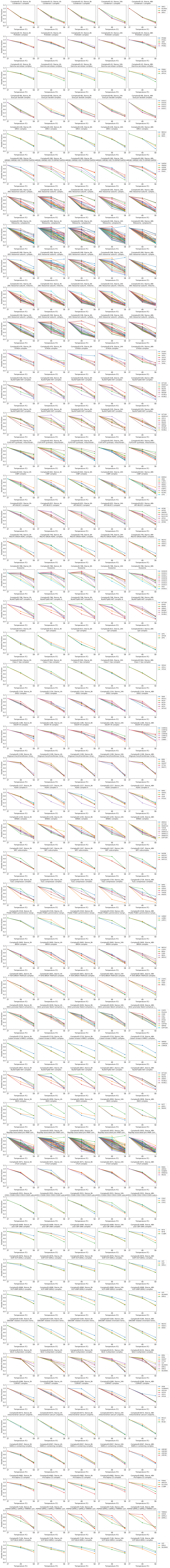

### Fig.S9_24h_convergent.pdf

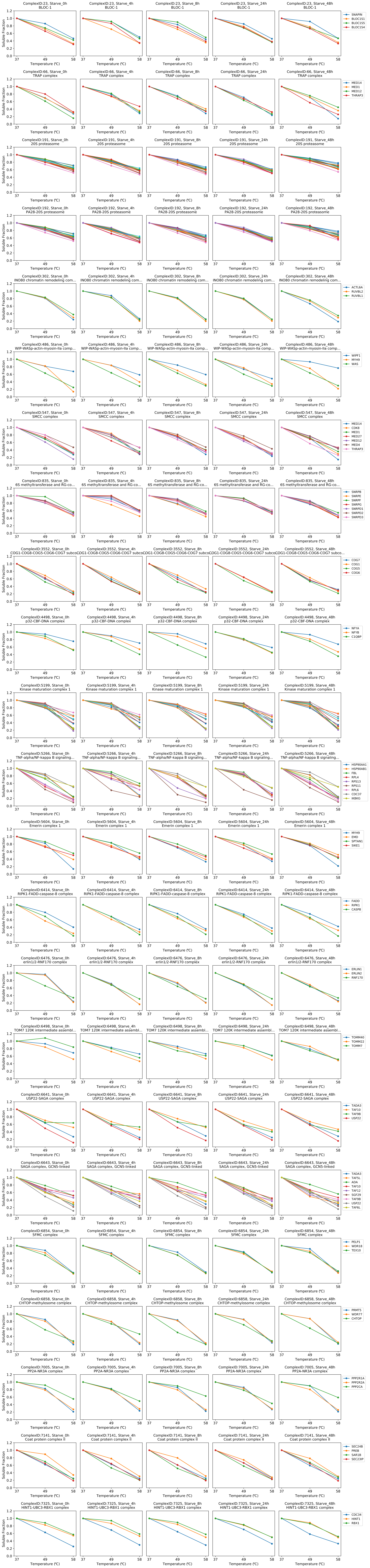

### Fig.S10_24h_divergent.pdf

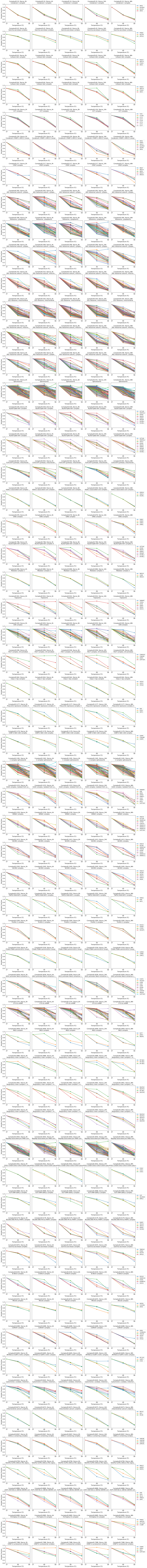

### Fig.S11_48h_convergent.pdf

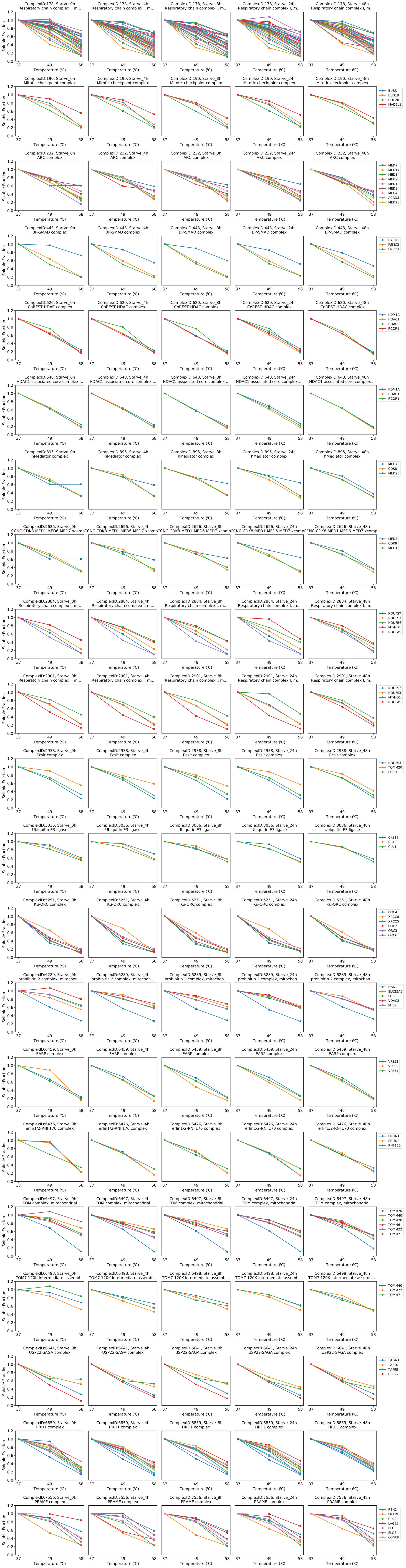

### Fig.S12_48h_divergent.pdf

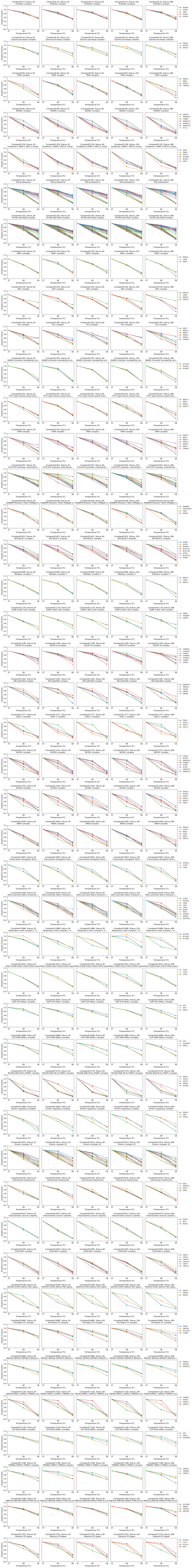

### Fig.S13_FACS_ayalysis.pdf

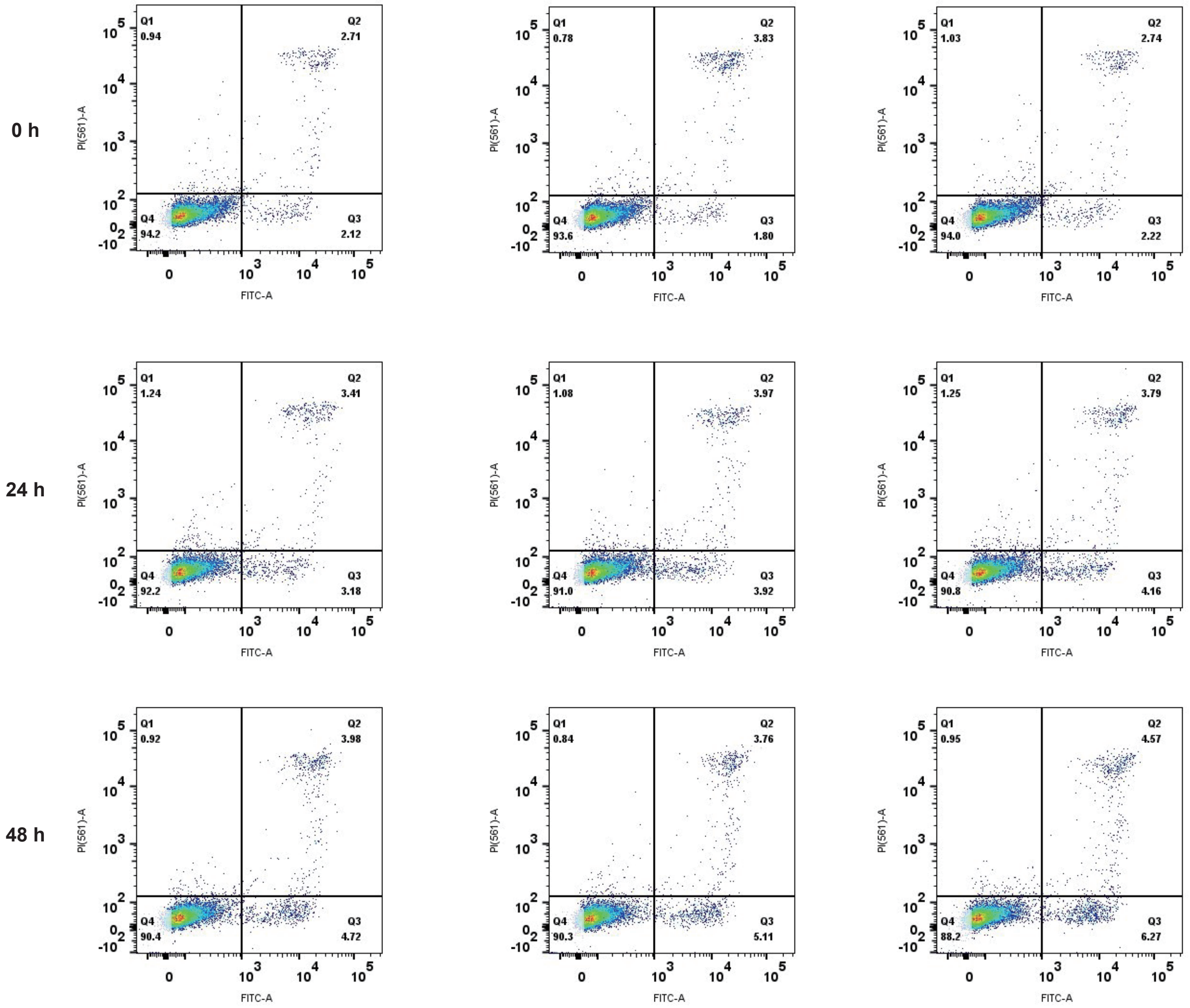

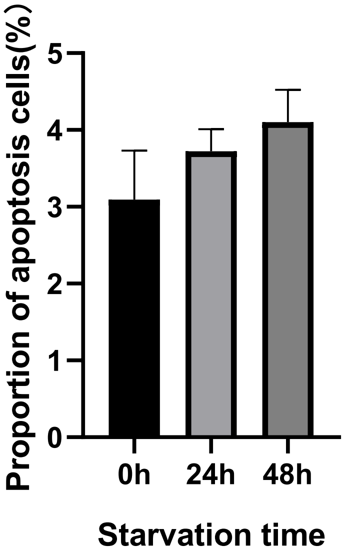

### Fig.S14_Corr_49.pdf

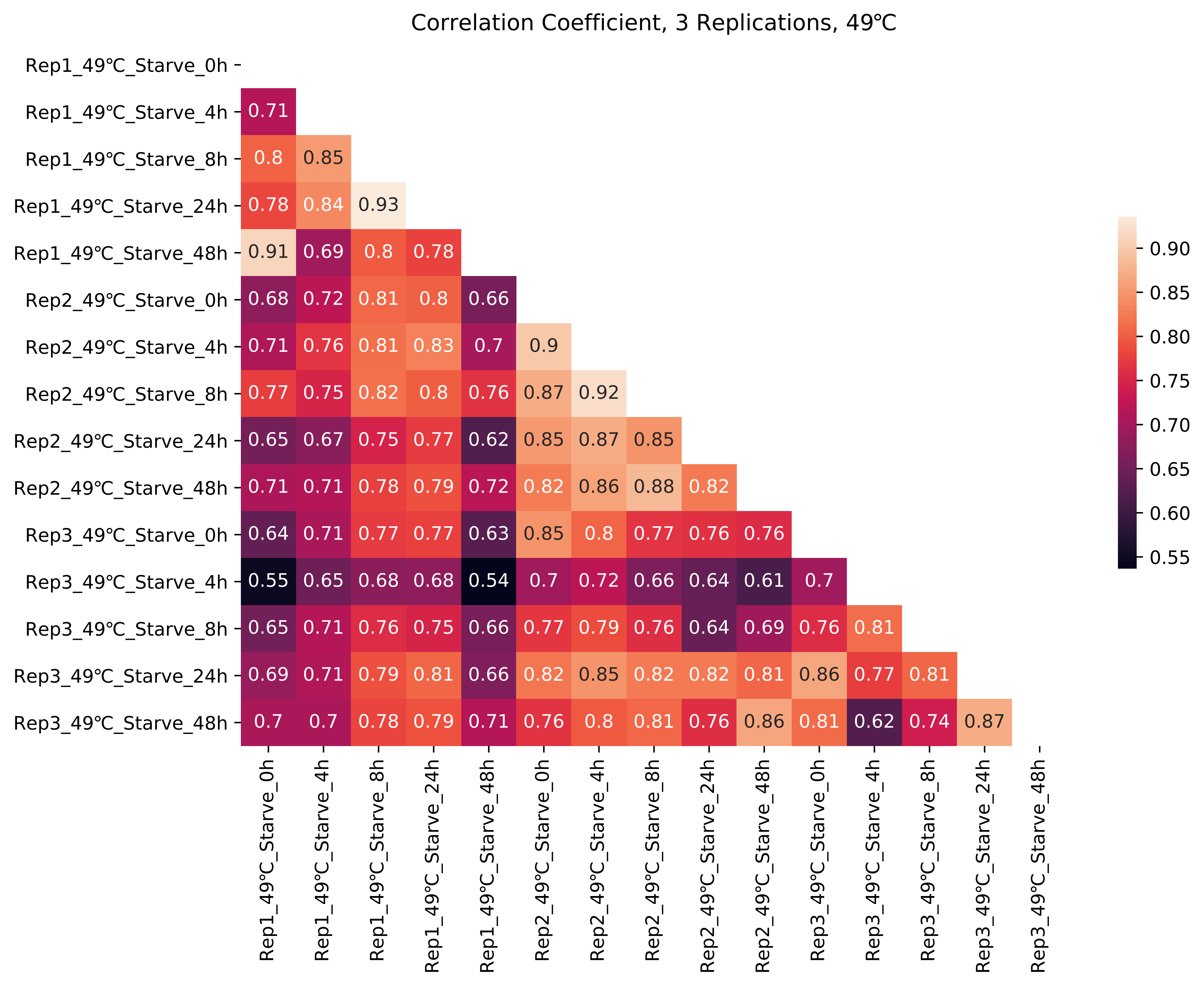

### Fig.S15_Corr_58.pdf

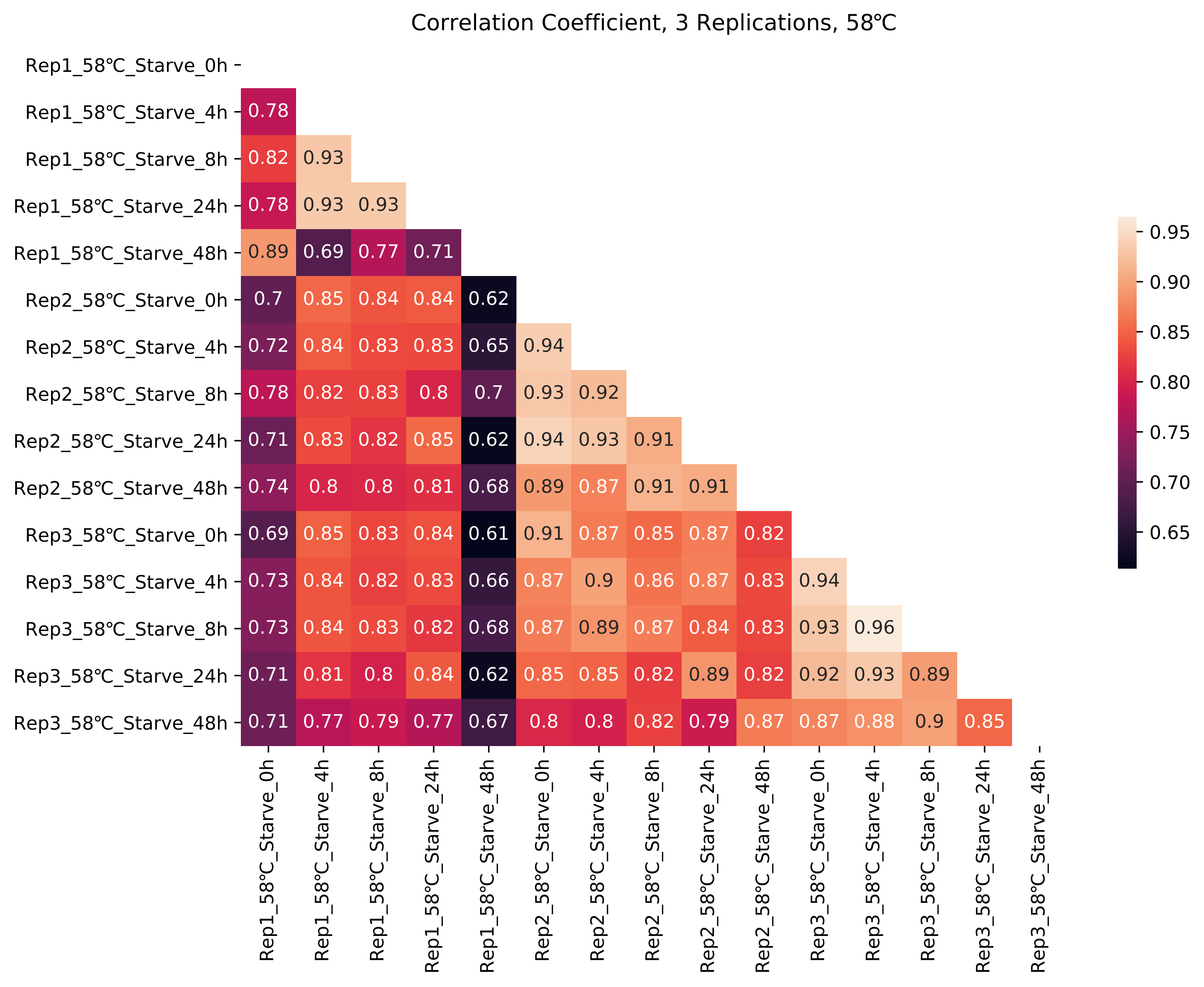

### Fig.S16.pdf

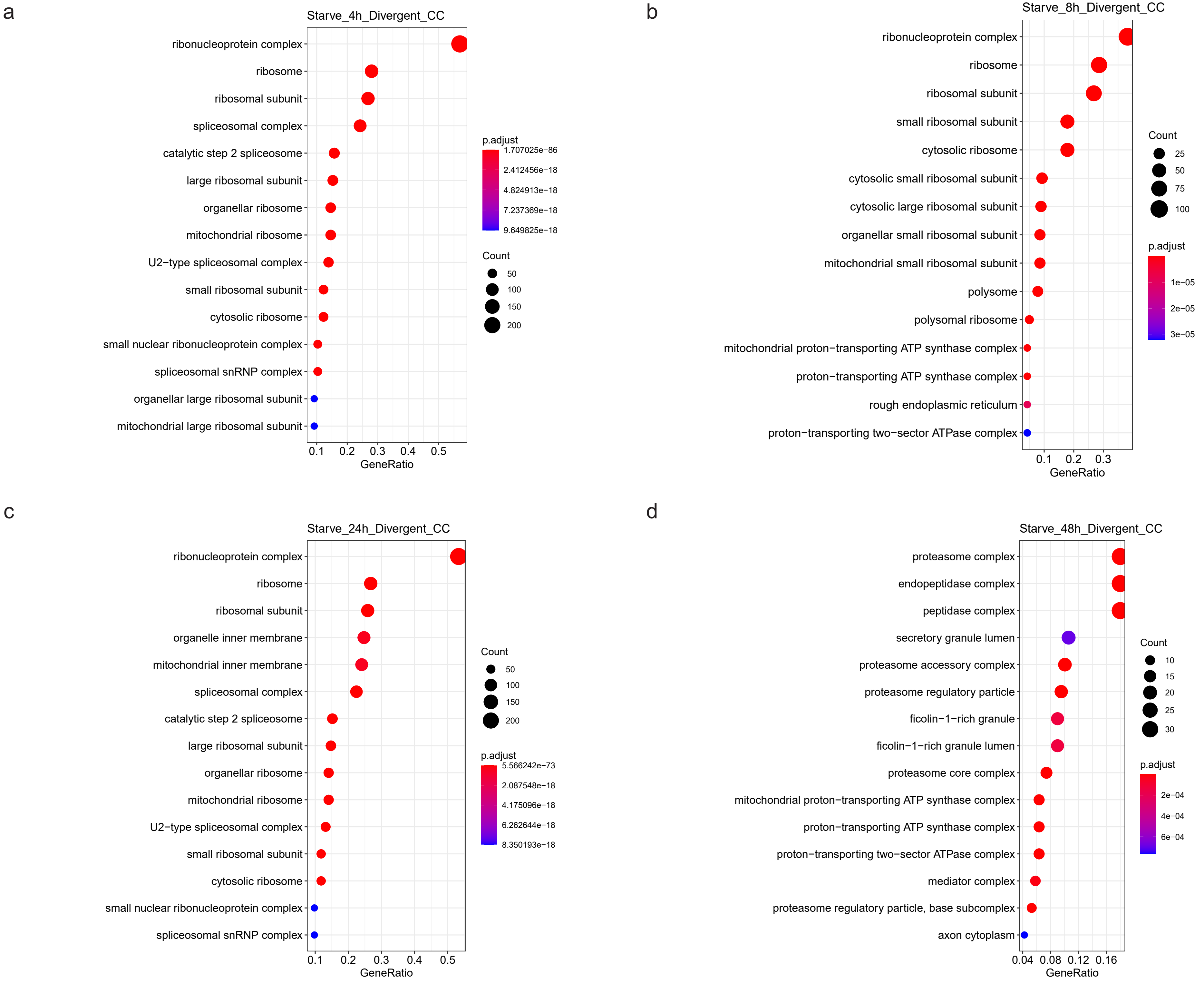

### Fig.S16a_GO_4h_divergent.pdf

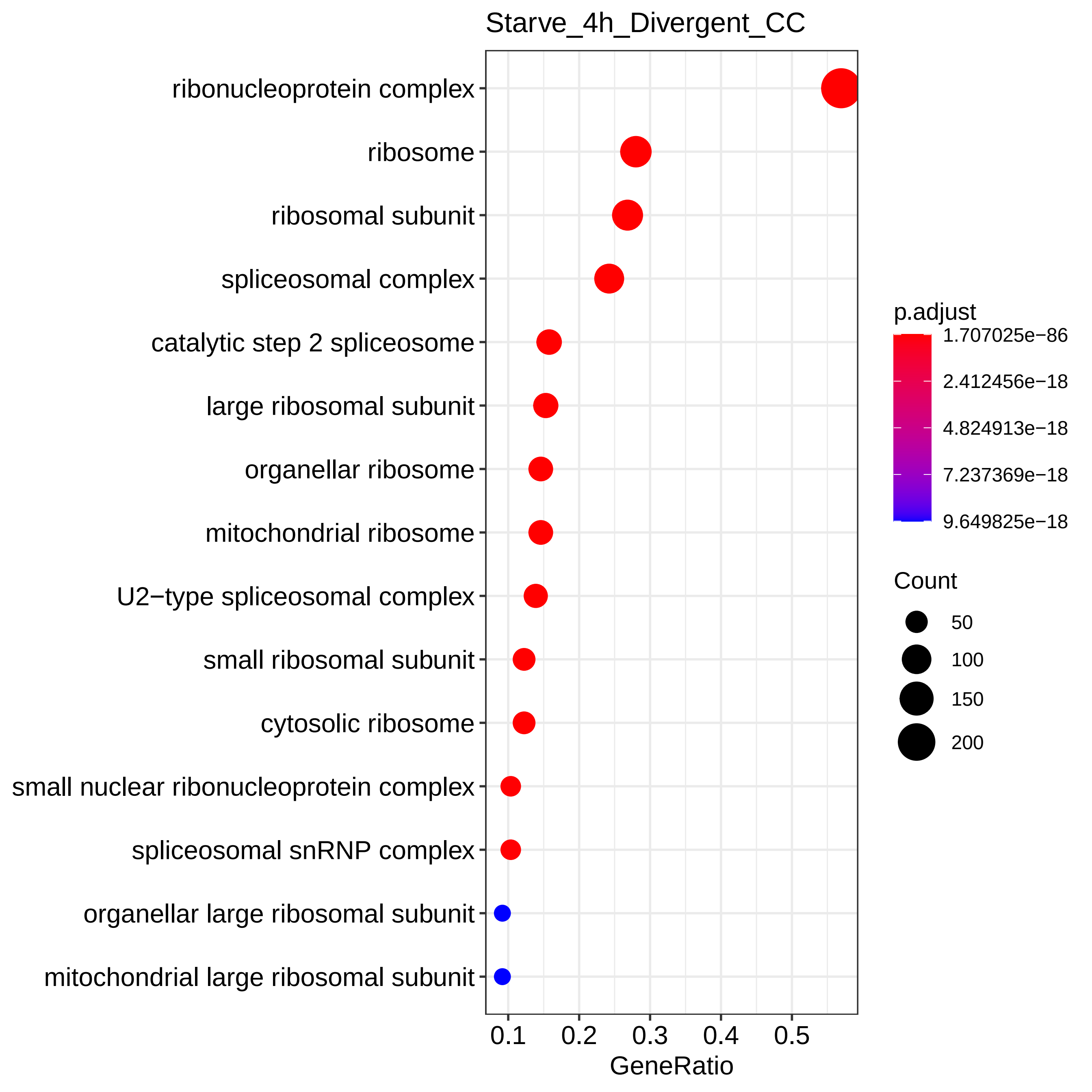

### Fig.S16b_GO_8h_divergent.pdf

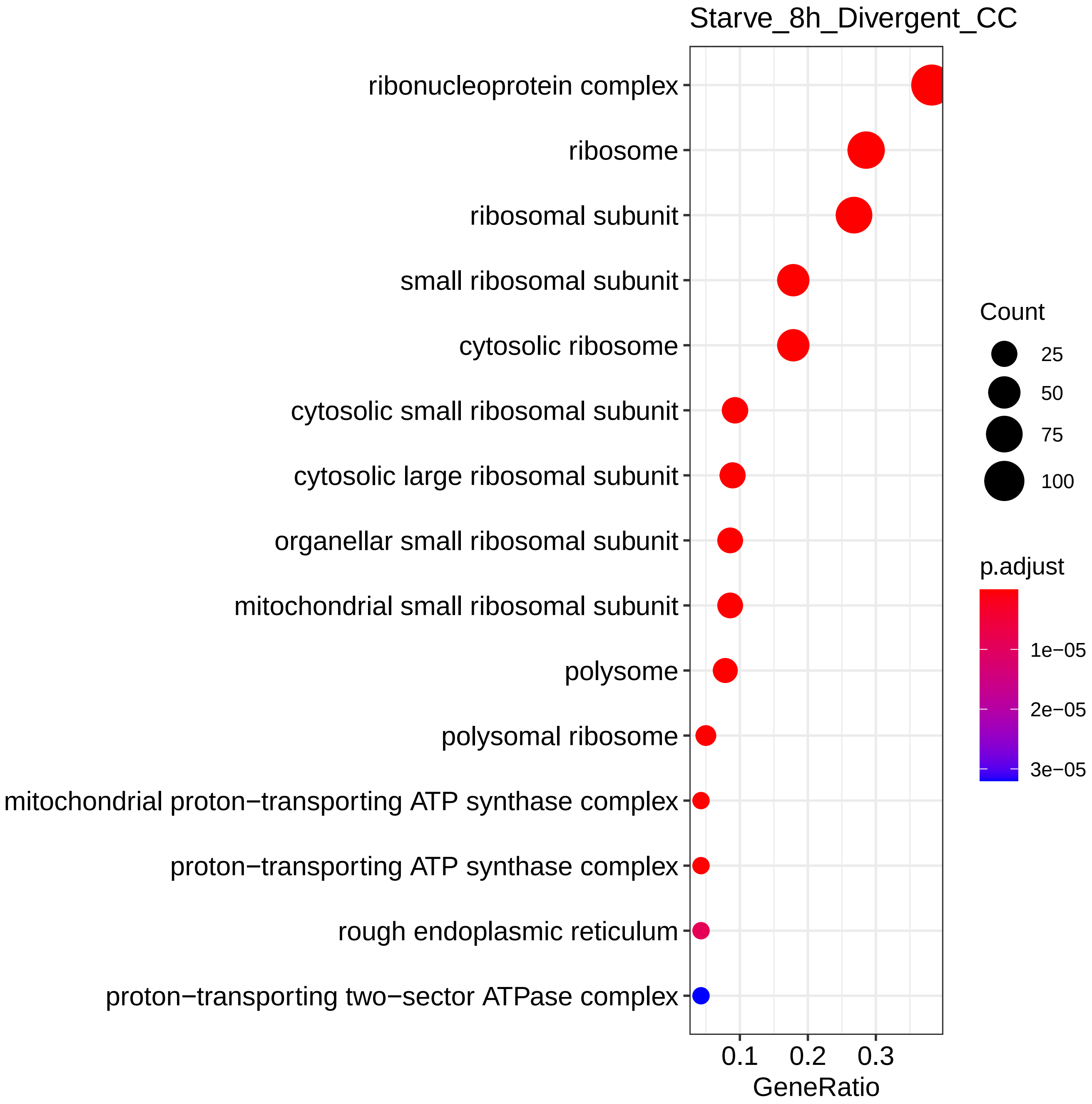

### Fig.S16c_GO_24h_divergent.pdf

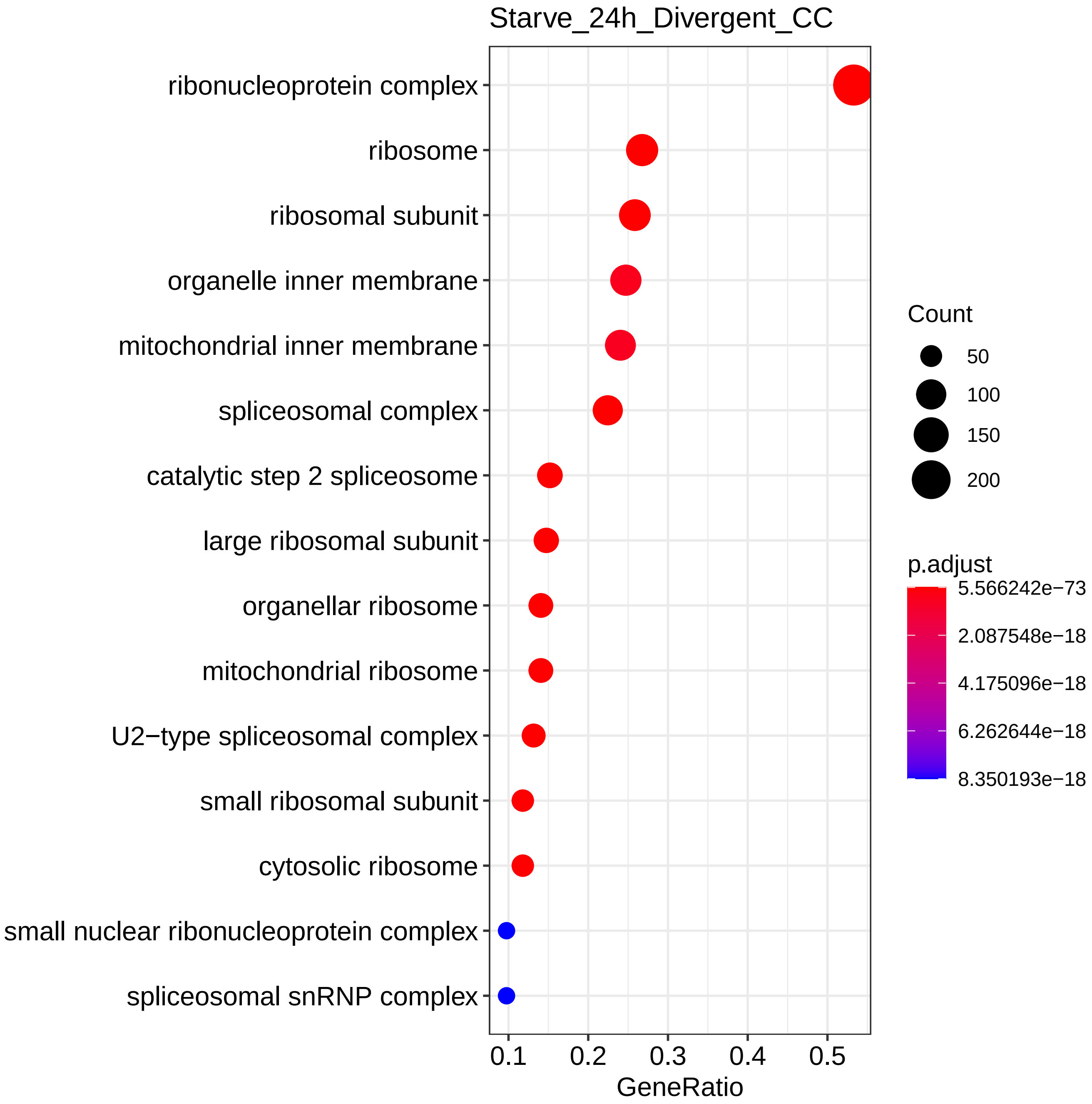

### Fig.S16d_GO_48h_divergent.pdf

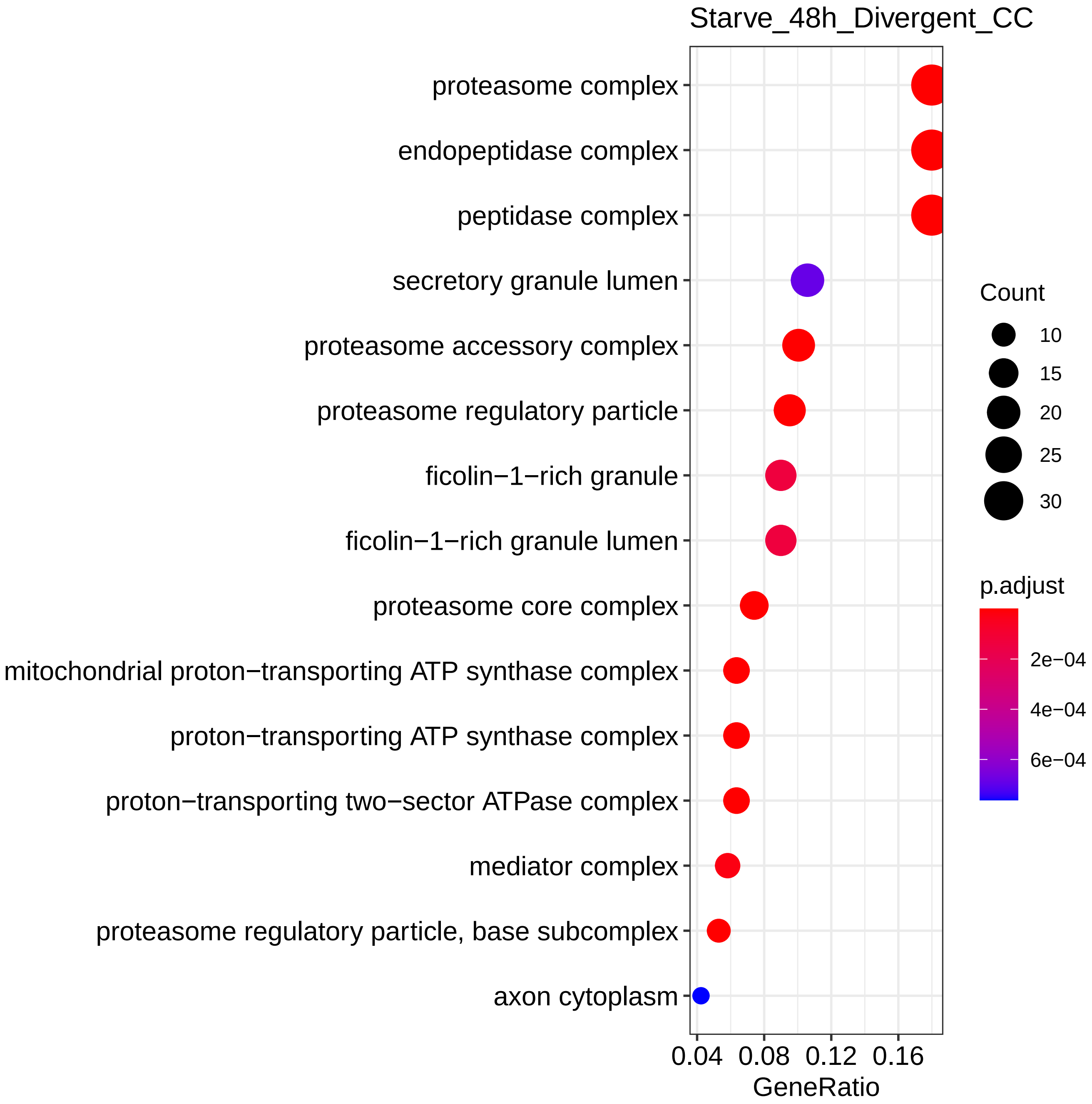
